# A General Birth-Death-Sampling Model for Epidemiology and Macroevolution

**DOI:** 10.1101/2020.10.10.334383

**Authors:** Ailene MacPherson, Stilianos Louca, Angela McLaughlin, Jeffrey B. Joy, Matthew W. Pennell

**Affiliations:** Department of Zoology and Biodiversity Research Centre, University of British Columbia, Vancouver, Canada; Department of Ecology and Evolutionary Biology, University of Toronto,Toronto, Canada; Department of Biology, University of Oregon, USA; Institute of Ecology and Evolution, University of Oregon, USA; British Columbia Centre for Excellence in HIV/AIDS, Vancouver, Canada; Bioinformatics, University of British Columbia, Vancouver, Canada; Department of Medicine, University of British Columbia, Vancouver, Canada

**Keywords:** epidemiology, macroevolution, phylogenetics, birth-death processes, statistical inference

## Abstract

Birth-death stochastic processes are the foundation of many phylogenetic models and are widely used to make inferences about epidemiological and macroevolutionary dynamics. There are a large number of birth-death model variants that have been developed; these impose different assumptions about the temporal dynamics of the parameters and about the sampling process. As each of these variants was individually derived, it has been difficult to understand the relationships between them as well as their precise biological and mathematical assumptions. Without a common mathematical foundation, deriving new models is non-trivial. Here we unify these models into a single framework, prove that many previously developed epidemiological and macroevolutionary models are all special cases of a more general model, and illustrate the connections between these variants. This framework centers around a technique for deriving likelihood functions for arbitrarily complex birth-death(-sampling) models that will allow researchers to explore a wider array of scenarios than was previously possible. We then use this framework to derive general model likelihoods for both the “single-type” case in which all lineages diversify according to the same process and the “multi-type” case, where there is variation in the process among lineages. By re-deriving existing single-type birth-death sampling models we clarify and synthesize the range of explicit and implicit assumptions made by these models.

---

As a consequence of their rapid mutation rates and large population sizes, many viral pathogens, such as HIV and SARS-CoV-2, accumulate genetic diversity on the timescale of transmission (Drummond et al., 2003; Duffy et al., 2008). This genetic diversity can be used to reconstruct the evolutionary relationships between viral variants sampled from different hosts, which in turn can help elucidate the epidemiological dynamics of a pathogen over time. The combined dynamics of viral diversification and the epidemiological process have been termed “phylodynamics” (Grenfell et al., 2004; Volz, 2012). In the last two decades, there has been a tremendous amount of innovation in phylodynamic methods, and the epidemiological inferences from these models increasingly complement those from more conventional surveillance data (Volz, 2012).

Phylodynamic models can be broadly grouped into two classes. The first, based on Kingman’s coalescent process (Kingman, 1982), has been historically widely used to examine changes in the historical population size of pathogens (Pybus et al., 2000; Strimmer and Pybus, 2001; Drummond et al., 2005; Volz et al., 2009). While appropriate in some applications, using the coalescent process depends upon the critical assumption that the population size is large relative to the number of samples, such that stochastic variation can be ignored; thus, this approach is inaccurate for reconstructing the dynamics of well-sampled or emerging pathogens (Stadler et al., 2015; Boskova et al., 2014). The second class of models, based on the birth-death process (Kendall, 1948; Maddison et al., 2007; Stadler, 2009, 2010), makes no assumptions about sparse sampling and fully incorporates stochasticity, and are thus become an increasingly favorable and popular alternative to coalescent models. These birth-death-sampling (BDS) models which are the focus of the present contribution, are also widely utilized in macroevolution to infer speciation and extinction rates over time (Raup, 1985; Nee et al., 1994; Morlon, 2014; Louca, 2020) and to estimate divergence times from phylogenetic data (Gernhard, 2008; Heath et al., 2014).

In the context of phylodynamics, BDS models have the additional property that the model parameters, which can be estimated from viral sequence data, explicitly correspond to parameters in classic structured epidemiological models that are often fit to case surveillance data. As the name implies, the BDS process includes three types of events: birth (pathogen transmission between hosts, or speciation in a macroevolutionary context), death (host death or recovery, or extinction in macroevolution), and sampling (including fossil collection in macroevolution). Taken together, these dynamics can be used to describe changes in the basic and effective reproductive ratios (*R*_0_ and *R*_*e*_, respectively) over time (Stadler et al., 2012, 2013) (see Box 1). A common inferential goal is to describe how the frequency of these events, and other derived variables such as *R*_*e*_, change throughout the course of an epidemic.

As we detail below and in the Supplementary Material, there has been an astounding rise in the variety and complexity of BDS model variants. A key assumption in the specification of BDS sub-models is whether all lineages are identical, diversifying according to the same process, or if rather the diversification process itself evolves (e.g., Maddison et al., 2007; FitzJohn, 2012; Stadler and Bonhoeffer, 2013; Rasmussen and Stadler, 2019; Barido-Sottani et al., 2018), with lineages belonging to one of multiple possible states each characterized by a unique process. As exemplified particularly by the range of available “single-type” models, each of these diversification processes can then be characterized by different dynamical assumptions. In the epidemiological case these assumptions specify, for example, the nature of viral transmission and the sampling procedure (Stadler et al., 2013; Kühnert et al., 2014; Gavryushkina et al., 2014). While typically not explicitly tied to mechanistic evolutionary processes, there are a similar abundance of dynamical assumptions employed in the macroevolutionary context specifying the nature of biodiversity change through time (Nee, 2006; Gernhard, 2008; Morlon et al., 2011; Stadler, 2011; Morlon, 2014; Heath et al., 2014; Louca, 2020).

This flourishing of methods and models has facilitated critical insights into epidemics (du Plessis and Stadler, 2015; Joy et al., 2016) and the origins of contemporary biodiversity (Morlon, 2014; Schluter and Pennell, 2017). However, this diversity of models has made it difficult to trace the connections between variants and to understand the precise epidemiological, evolutionary, and sampling processes that differ between them. Furthermore, despite their apparent similarities, these models have been derived on a case-by-case basis using different notation and techniques; this creates a substantial barrier for researchers working to develop novel models for new situations. And critically, it is imperative that we understand the general properties of BDS phylogenetic models and the limits of inferences from them (Louca and Pennell, 2020a; Louca et al., 2021) and this is difficult to do without considering the full breadth of possible scenarios.

Here we address all of these challenges by providing a unified framework for deriving the probability of observing a given phylogeny and series of sampling events under a particular BDS model. We demonstrate the utility of five-step procedural framework to derive both a general single-type and a general multi-type model likelihood. Specifically, these general models do not assume anything about the functional forms (i.e., temporal dynamics) of the various parameters including sampling, the possibility of sampling ancestors (or not), or how the process was conditioned. Such general models may be useful for studying the mathematical properties of sets of models (Louca and Pennell, 2020a; Louca et al., 2021). However, as they are computationally cumbersome these general models will not necessarily be useful for statistical inference. Rather the procedural framework and the general model likelihood can streamline derivation of new sub-model variants adapted to specific applications.

In addition to the derivation of the general models, in the single-type case we use the procedural framework to re-derive a wide array of existing BDS models, clarifying the assumptions of each and providing likelihood expressions with standardized notation. In the multi-type case, this framework builds on recent work (Barido-Sottani et al., 2020) to provide an alternative formulation of many of the existing model likelihoods. We then use the specific case of the BiSSEness model (Magnuson-Ford and Otto, 2012; Goldberg and Igić, 2012) to illustrate how the tree likelihood can be calculated either through the traditional post-order traversal algorithm or via this five-step framework. Given our summary of single-type BDS models and the well known connections among such multi-type BDS models (for recent reviews see Morlon, 2014; Ng and Smith, 2014), our BDS framework hence provides a unification and extension to an extensive class of methods employed across epidemiological and macroevolutionary contexts.

## The single-type birth-death-sampling model

### Model Specification

The BDS stochastic process begins with a single lineage at time *T* before the present day. We note that this may be considerably older than the age of the most recent common ancestor of an observed sample which is given by *t*_MRCA_. While we focus primarily on applications to epidemiology, our approach is agnostic to whether the rates are interpreted as describing pathogen transmission or macroevolutionary diversification.

In the model, transmission/speciation results in the birth of a lineage and occurs at rate *λ*(*τ*), where *τ* (0 ≤ *τ ≤ T*) is measured in units of time before the present day, such that *λ* can be time-dependent. We make the common assumption that lineages in the viral phylogeny coalesce exactly at transmission events, thus ignoring the pre-transmission interval inferred in a joint phylogeny of within- and between-host sequences (Romero-Severson et al., 2016). Throughout, we will use *τ* as a a general time variable and *t*_×_ to denote a specific time of an event × units before the present day (see Table S1). Lineage extinction, resulting from host recovery or death in the epidemiological case or the death of all individuals in a population in the macroevolutionary case, occurs at time-dependent rate *μ*(*τ*). We allow for two distinct types of sampling: lineages are either sampled according to a Poisson process through time *ψ*(*τ*) or binomially at very short intervals, which we term “concerted sampling attempts” (CSAs), where lineages at some specified time *t*_*l*_ are sampled with probability *ρ*_*l*_ (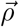 denotes a vector of concerted sampling events at different time points). In macroevolutionary studies based only on extant lineages, there is no Poissonian sampling, but a CSA at the present (*ρ*_0_ *>* 0). In epidemiology, CSAs correspond to large-scale testing efforts (relative to the background rate of testing) in a short amount of time (relative to the rates of viral sequence divergence); for full explanation, see Appendix. We call these attempts rather than events because if *ρ* is small or the infection is rare in the population, few or no samples may be obtained. CSAs can also be incorporated into the model by including infinitesimally short spikes in the sampling rate *ψ* (more precisely, appropriately scaled Dirac distributions). Hence, for simplicity, in the main text we focus on the seemingly simpler case of pure Poissonian sampling through time except at present-day, where we allow for a CSA to facilitate comparisons with macroevolutionary models; the resulting formulas can then be used to derive a likelihood formula for the case where past CSAs are included (see Appendix).

In the epidemiological case, sampling may be concurrent (or not) with host treatment or behavioural changes resulting in the effective extinction of the viral lineage. Hence, we assume that sampling results in the immediate extinction of the lineage with probability *r*(*τ*). As with the CSAs, this arbitrary time dependence allows for the incorporation of Dirac spikes in any of these variables, for example with mass extinctions (*μ*) and lagerstätten in the fossil record (*ψ*(1 − *r*)) (Magee and Höhna, 2021). Similarly, in the case of past CSAs we must include the probability, *r*_*l*_, that sampled hosts are removed from the infectious pool during the CSA at time *t*_*l*_. Poissonian sampling without the removal of lineages (*r*(*τ*) < 1) can be employed in the macroevolutionary case to explicitly model the collection of samples from the fossil record (such as the fossilized-birth-death process; Heath et al., 2014).

For our derivation, we make no assumption about the temporal dynamics of *λ*, *μ*, *ψ*, or *r*; each may be constant, or vary according to any arbitrary function of time given that it is biologically valid (non-negative and between 0 and 1 in the case of *r*). We make the standard assumption that at any given time any given lineage experiences a birth, death or sampling event independently of (and with the same probabilities as) all other lineages. We revisit this assumption in Box 1 where we discus how the implicit assumptions of the single-type BDS process are well summarized by the diversification model’s relationship to the SIR epidemiological model. Our resulting general time-variable BDS process can be fully defined by the parameter set 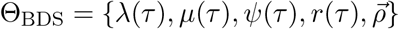.

In order to make inference about the model parameters, we need to calculate the likelihood, 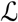, that an observed phylogeny, 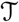, is the result of a given BDS process. With respect to the BDS process there are two ways to represent the information contained in the phylogeny 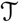, both of which have been used in the literature, which we call the “edge” and “critical time” representations, respectively. We begin by deriving the likelihood in terms of the edge representation and later demonstrate how to reformulate the likelihood in terms of critical times. In the edge representation, the phylogeny is summarized as a set of edges in the mathematical graph that makes up the phylogeny, numbered 1-11 in Figure B1 panel C, and the types of events that occurred at each node. We define *g*_*e*_(*τ*) as the probability that the edge *e* which begins at time *s*_*e*_ and ends at time *t*_*e*_ gives rise to the subsequently observed phylogeny between time *τ,* (*s*_*e*_ < τ < *t*_*e*_) and the present day. The likelihood of the tree then, is by definition *g*_*stem*_(*T*): the probability density the stem lineage (stem = 1 in Figure B1 panel C) gives rise to the observed phylogeny from the origin, *T*, to the present day. Although it is initially most intuitive to derive the likelihood in terms of the edge representation, as we show below, it is then straightforward to derive the critical times formulation which results in mathematical simplification.

### 1. Deriving the Initial Value Problem (IVP) for g_e_(τ)

We derive the IVP for the likelihood density *g*_*e*_(*τ*) using an approach first developed by Maddison et al. (2007). For a small time Δ*τ* the recursion equation for the change in the likelihood density is given by the following expression.

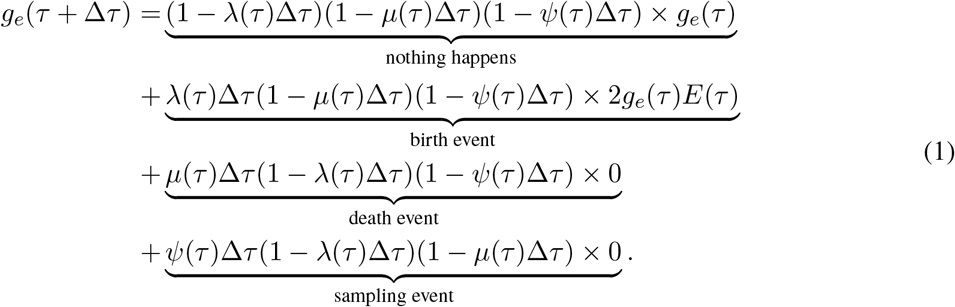

Here, *E*(*τ*) is the probability that a lineage alive at time *τ* leaves no sampled descendants at the present day. We will examine this probability in more detail below. Assuming Δ*τ* is small, we can approximate the above recursion equation as the following difference equation.

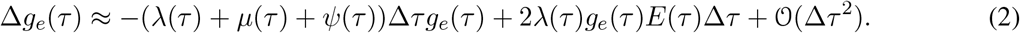

By the definition of the derivative we have:

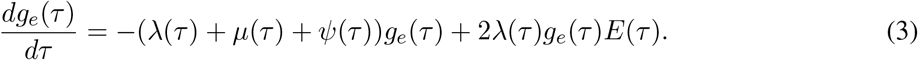

Equation (3) is known as the Kolmogorov backward equation of the BDS process (Feller, 1949; Louca and Pennell, 2020b). Beginning at time *s*_*e*_, the initial condition of *g*_*e*_ depends on which event occurred at the beginning of edge *e*.

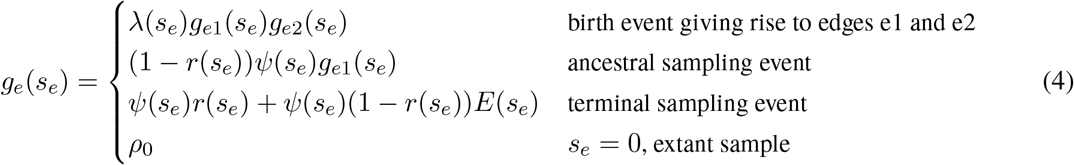

Together Equations (3) and (4) define the initial value problem for *g*_*e*_(*τ*) as a function of the probability *E*(*τ*).

Because the likelihood density *g*_*e*_ is the solution to a linear differential equation with initial condition at time *s*_*e*_, we can express its solution as follows:

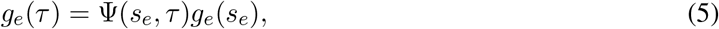

where the auxiliary function, Ψ, is given by:

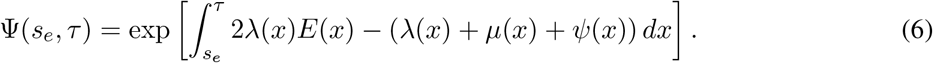

This function, Ψ(*s, t*), maps the value of *g*_*e*_ at time *s* to its value at *t*, and hence is known as the probability “flow” of the Kolmogorov backward equation (Louca and Pennell, 2020b).

### 2. Deriving the IVP for E(τ)

We derive the IVP for *E*(*τ*) in a similar manner as above, beginning with a difference equation.

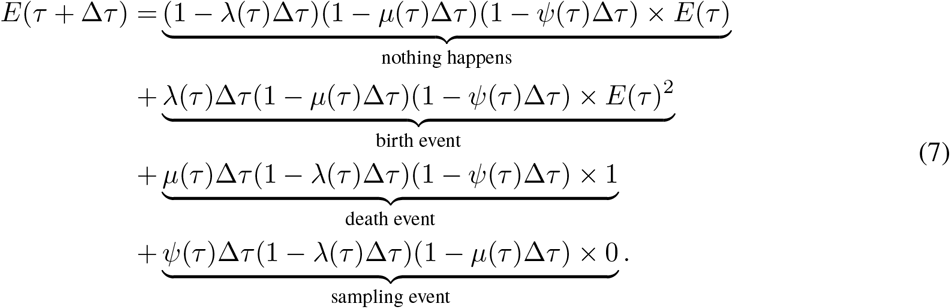

By the definition of a derivative we have:

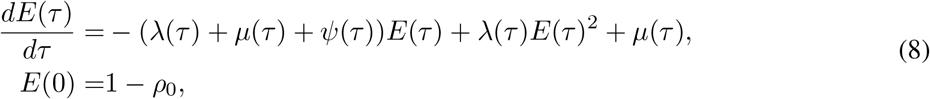

where *ρ*_0_ is the probability a lineage is sampled at the present day. The initial condition at time 0 is therefore the probability that a lineage alive at the present day is not sampled. Given an analytical or numerical general solution to *E*(*τ*), we can find the likelihood by evaluating *g*_*stem*_(*T*), as follows.

### 3. Deriving the expression for g_stem_(T)

Given the linear nature of the differential equation for *g*_*e*_(*τ*) and hence the representation in Equation (5)), the likelihood *g*_stem_(*τ*) is given by the product over all the initial conditions times the product over the probability flow for each edge.

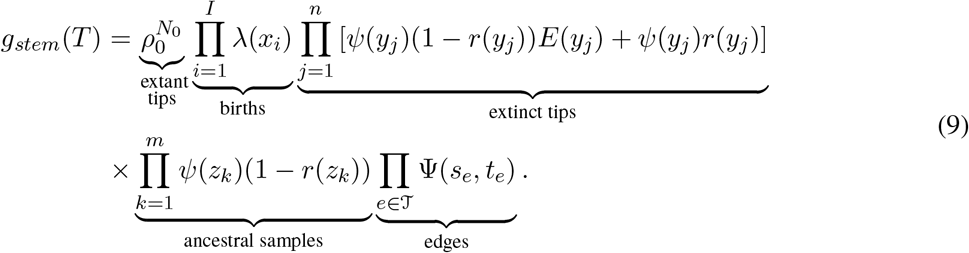

### 4. Representing g_stem_(T) in terms of critical times

Equation (9) can be further simplified by removing the need to enumerate over all the edges of the phylogeny (the last term of Equation (9)) and writing 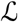 in terms of the tree’s critical times (horizontal lines in figure B1). The critical times of the tree are made up of three vectors, 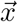, 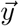, and 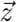, as well as the time of origin *T*. The vector 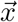 gives the time of each birth event in the phylogeny and has length *I* = *N*_0_ − *n* − 1 where *N*_0_ is the number of lineages sampled at the present day and *n* is the number of terminal samples. Unless noted otherwise the elements of vector 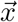 are listed in decreasing order, such that *x*_1_ *> x*_2_ *> ...x*_*I*_ and hence *x*_1_ is the time of the most recent common ancestor *t*_MRCA_. The vector 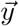 gives the timing of each terminal sample and hence has length *n* whereas vector 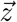 gives the timing of each ancestral sample and has length *m*. With respect to the BDS likelihood then the sampled tree is summarized by 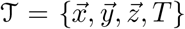. We note that the critical times only contain the same information as the edges as a result of the assumptions of the BDS process but are not generally equivalent representations of 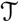.

As a result of the linear nature of *g*_*e*_(*τ*) it is straightforward to rewrite the likelihood in Equation (9) in terms of the critical-time representation of the sampled tree. Defining

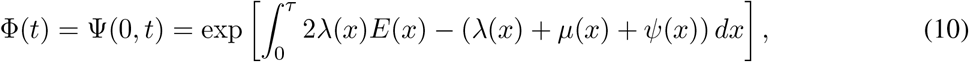

the probability flow Ψ can be rewritten as the following ratio:

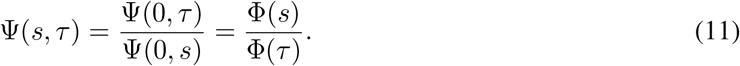

This relationship allows us to rewrite the likelihood by expressing the product over the edges as two separate products, one over the start of each edge and the other over the end of each edge. Edges begin (value of *t*_*e*_) at either: 1) the tree origin, 2) a birth event resulting to two lineages, or 3) an ancestral sampling event. Edges end (values of *s*_*e*_) at either: 1) a birth event, 2) an ancestral sampling event, 3) a terminal sampling event, or 4) the present day. Hence we have:

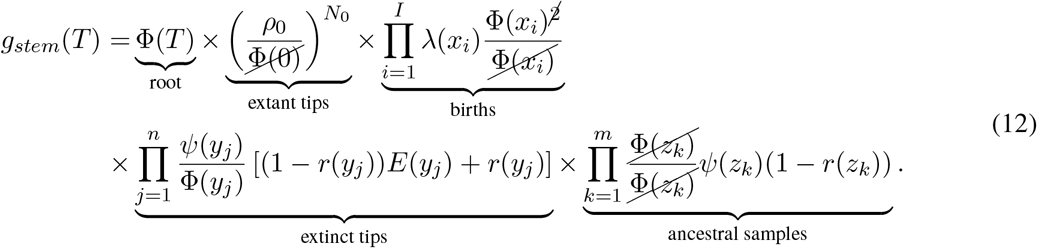

Note Φ(0) = 1.

### 5. Conditioning the likelihood

While Equation (12) is equal to the basic likelihood of the phylogeny 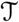, it is often appropriate to condition the tree likelihood on the tree exhibiting some property, for example the condition there being at least sampled lineage. Imposing a condition on the likelihood is done by multiplying by a factor 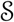. Various conditioning schemes are considered in section 1 and listed in Table S3. The resulting likelihood of the general BDS model is:

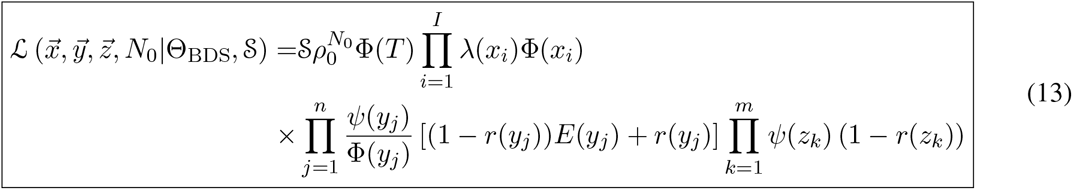

## Many existing models are special cases of this general BDS model

A large variety of previously published BDS models in epidemiology and macroevolution are simply special cases of the general model presented here (for a summary of the models we investigated see Table S2; proofs in Supplemental Material). Indeed, we can obtain the likelihood of these models by adding mathematical constraints (i.e., simplifying assumptions) to the terms in Equation (13). Our work thus not only provides a consistent notation for unifying a multitude of seemingly disparate models, it also provides a concrete and numerically straightforward recipe for computing their likelihood functions. We have implemented the single-type BDS likelihood in the R package `castor` (Louca and Doebeli, 2018), including routines for maximum-likelihood fitting of BDS models with arbitrary functional forms of the parameters given a phylogeny and routines for simulating phylogenies under the general BDS models (functions `fit_hbds_model_on_grid, fit_hbds_model_parametric and generate_tree_hbds`).

Figure 1 summarizes the simplifying assumptions that underlie common previously published BDS models; these assumptions generally fall into four categories: 1) assumptions about the functional form of birth, death, and sampling rates over time, 2) assumptions pertaining to the sampling of lineages, 3) the presence of mass-extinction events, and 4) the nature of the tree-conditioning as given by 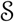. Here we provide a brief overview of the type of previously-invoked constraints which are consistent (or not) with our unified framework; for full details on each specific case, we refer readers to the Supplementary Material. While we illustrate these constraints within the single-type context, analogous assumptions can be made within the multi-type context examined in the following section. In regards to rate assumptions, many early BDS models (Stadler, 2009, 2010; Stadler et al., 2012) assumed that the birth, death, and sampling rates remained constant over time. This is mathematically and computationally convenient since an analytical solution can easily be obtained for *E*(*τ*). In the epidemiological case, holding *λ* constant, however, implies that the number of susceptible hosts is effectively constant throughout the epidemic and/or that the population does not change its behavior over time (an unrealistic assumption given seasonal changes or changes in response to the disease itself). As such, this assumption is only really valid for small time periods or the early stages of an epidemic. This is useful for estimating the basic reproductive number, *R*_0_, of the SIR model (Box 1) but not for the effective reproductive number *R*_*e*_ at later time points (Stadler et al., 2012).

**Figure 1:**
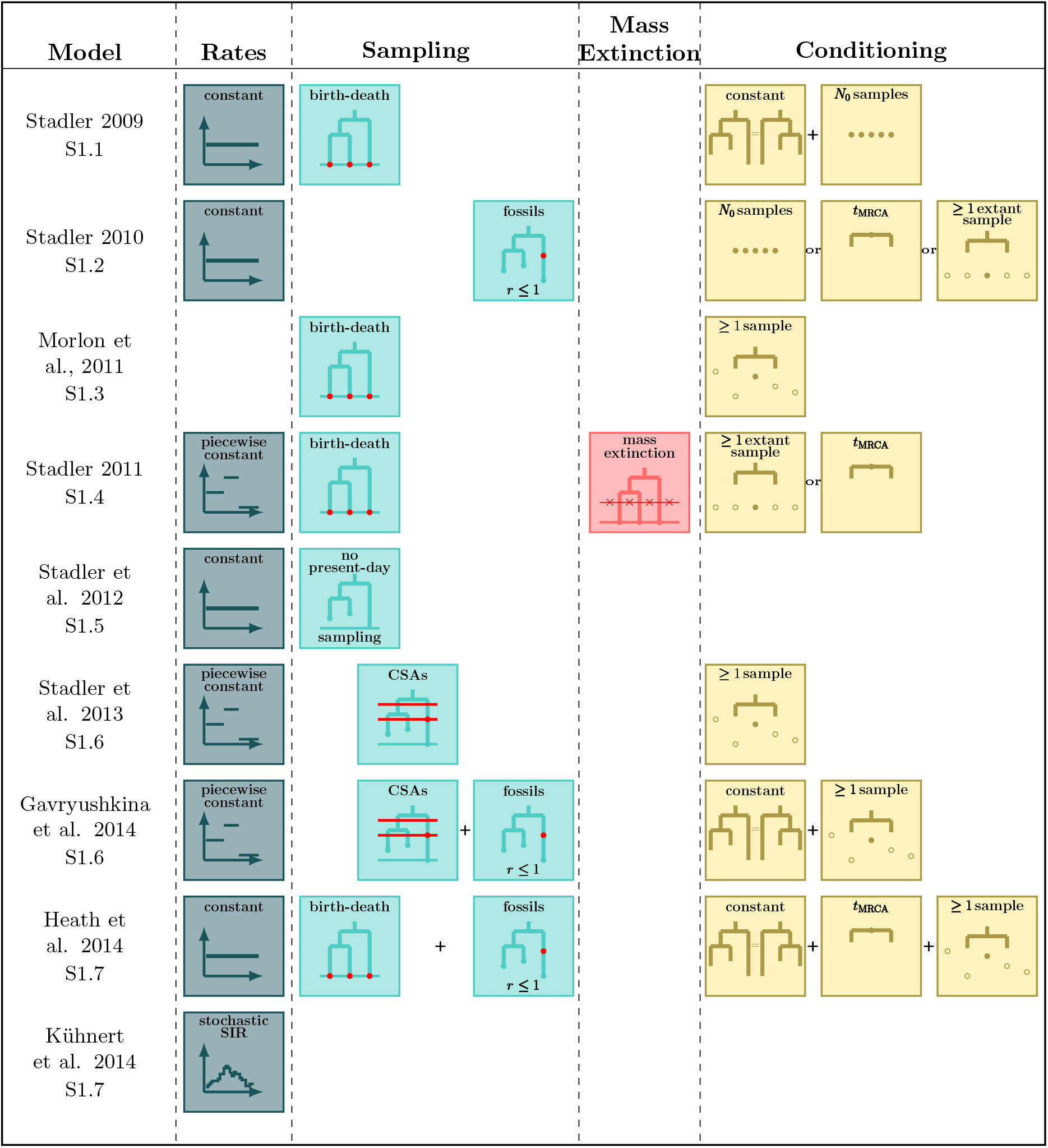
Sub-model assumptions. Rate, sampling, mass extinction, and conditioning assumptions of existing sub-models of the general time-variable BDS process. The key points are that i) each of the previously developed models we considered can be obtained by adding specific combinations of constraints to the various parameters of the general BDS model; and ii) that there are many plausible, and potentially biologically informative combinations of constraints that have not been considered by researchers in epidemiology or macroevolution.

A similarly tractable, but more epidemiologically relevant, model is known as the “birth-death-skyline” variant (Stadler and Bonhoeffer, 2013; Gavryushkina et al., 2014), in which rates are piecewise-constant functions through time (like the constant rate model, there is also an analytical way to calculate the likelihood of this model; see Appendix). The BDS skyline model has been implemented under a variety of additional assumptions in the Bayesian phylogenetics software BEAST Bouckaert et al. (2019). The BDS skyline model has also been extended by Kuhnert et al. (Kühnert et al., 2014) to infer the the parameters of an underlying stochastic SIR model. In this case the diversification model parameters Θ_*BDS*_ are random variables that emerge from stochastic realizations of the epidemiological model given by Θ_*SIR*_, see Equation (B1). Finally, the birth-death skyline model with piecewise constant rates can also be applied in the macroevolutionary case when no sampling occurs through time, *Ψ*(*τ*) = 0 (Stadler, 2011).

In addition to imposing constraints on the temporal variation in the rates, previously derived sub-models have considered a variety of different assumptions about the nature of the sampling process. Most notably, in macroevolutionary studies, sampling of molecular data typically occurs only at the present day (Stadler, 2009, 2011; Morlon et al., 2011) whereas past Poissonian sampling can be introduced to include the sampling of fossil data (Heath et al., 2014). In epidemiology, concerted sampling at the present day is likely biologically unrealistic (Stadler et al., 2012), though in some implementations of the models, such a sampling scheme has been imposed. These concerted sampling attempts prior to the present day as well as mass extinction events can be incorporated via the inclusion of Dirac distributions in the sampling and death rates, respectively. Finally, previous models often multiply the likelihood by a factor 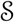 in order to condition on a particular observation (e.g., observing at least one lineage or exactly *N*_0_ lineages), enumerate of indistinguishable trees (Gavryushkina et al., 2013, 2014; Stadler, 2009), or to reflect known uncertainties. The “fossilized-birth-death” likelihood derived by Heath et al. (2014) for example, includes a factor that reflects the uncertainty in the attachment and placement of fossils on the macroevolutionary tree. This fossilized-birth-death process has been used to estimate divergence times and to model lineage diversification (Gavryushkina et al., 2017; Landis et al., 2021). Variants of the fossilized-birth-death process, for example including mass extinction events, are feasible and can be derived using our approach. We also note that models similar to the time-variable fossilized-birth-death process have been developed for cases when phylogenetic data is not available (i.e., when only including fossil occurrence data Silvestro et al., 2014; Lehtonen et al., 2017); we have not investigated how these models relate to our generalized BDS model but we speculate that it would be possible to also bring these models into a common framework with those that we have discussed. The Supplementary Material demonstrates how these sub-models can be re-derived by either imposing the necessary constraints on the general likelihood formula given in Equation (13) or, alternatively, by starting from the combinations of assumptions and using the five-step procedure outlined above.

## The multi-type birth-death-sampling model

A common extension of the single-type diversification models explored above is to consider cases where the diversification rates (*λ, μ, Ψ*) and probabilities (*r, ρ*) vary among lineages as a function of a categorical “lineage type”. This lineage type can be defined in terms of specific (e.g., Maddison et al., 2007; Rasmussen and Stadler, 2019) or unspecified traits (e.g., Beaulieu and O’Meara, 2016) or trait combinations (FitzJohn, 2012) (for recent reviews of these models see Morlon, 2014; Ng and Smith, 2014). Representing these lineage types as colours at nodes and along branches of the tree, we first extend the the single-type model above by deriving the likelihood of a fully coloured tree with topology 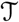 where the states along all edges of the phylogeny are known as given by 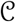. The resulting likelihood is an extension of the likelihood first developed by (Barido-Sottani et al., 2020), where the diversification rates and probabilities are allowed to vary arbitrarily through time. Following the approach of Magnuson-Ford and Otto (2012) and Goldberg and Igić (2012), the state of lineages can change either anagenetically, with a lineage of type *a* mutating to a type *b* at rate *γ*_*a,b*_(*τ*) or cladogenetically, with a lineage of type *a* giving rise to a daughter lineage of type *b* at rate *λ*_*a,b*_(*τ*). Lineages go extinct at a state-dependent rate *μ*_*a*_(*τ*) and are sampled at rate *Ψ*_*a*_(*τ*). As in the single-type model, upon sampling lineages are removed from the population with probability *r*_*a*_(*τ*) whereas all lineages alive at the present day are sampled with a probability *ρ*_*a*_(*τ*).

We use the five-step framework specified above for the single-type case to derive a general multi-type model likelihood of the coloured tree follows in the same manner as the single-type model (see supplementary material II). We first derive the initial value problem for the probability *g*_*e,a*_(*τ*) that an edge *e* of type *a* in the tree at time *τ* gives rise to the subsequently observed phylogeny. The edge *e* here refers not to an edge in the topological tree, but to a segment of the tree all of one state between birth, sampling, or mutation events.

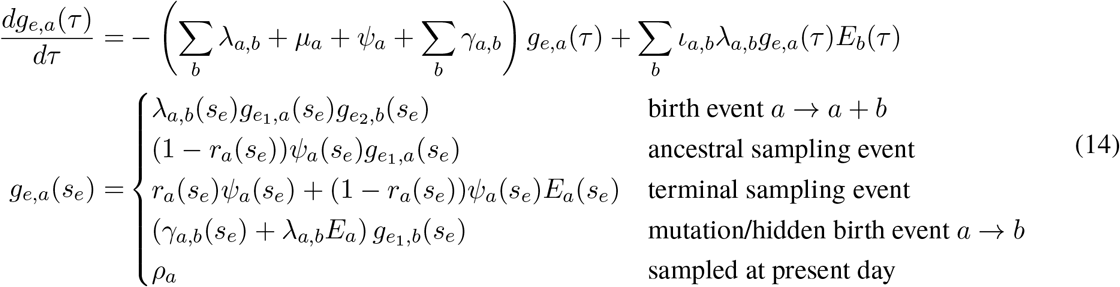

where *ι*_*a,b*_ is an indicator variable which takes on value of 2 if *a* = *b* and 1 otherwise. Importantly the differential equation for *g*_*e,a*_ is linear and hence has a known general solution *g*_*e,a*_ = *g*_*e,a*_(*s*_*e*_)Ψ(*s*_*e*_, *τ*). As in the single-type model Ψ(*s*_*e*_, *τ*) is the probability flow (Louca and Pennell, 2020b) mapping the probability *g*_*e,a*_ from the initial state at time *s*_*e*_ to the probability at time *τ*.

An analogous initial value problem can be derived for the probability *E*_*a*_(*τ*), that a lineage of type *a* alive at time *τ* leaves no observed descendants in the sampled tree.

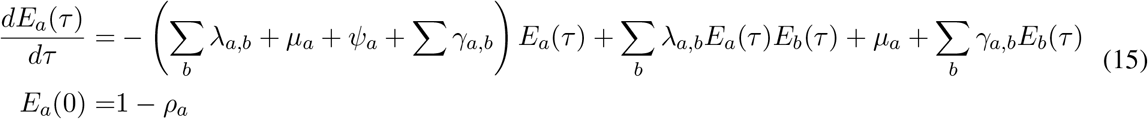

This is a non-linear differential equation and must be solved numerically. Given the solution of *g*_*e,a*_ and *E*_*a*_ the likelihood for the fully coloured tree is characterized by a series of critical times: first, 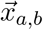 the times at which a lineage of type *a* gives birth to a lineage of type *b*, 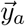 the ages of tip samples of type *a*, 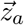 the ages of ancestral samples of type *a*, and 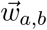 the times at which lineages are observed to transition events from type *a* to type *b*. The resulting likelihood is given by:

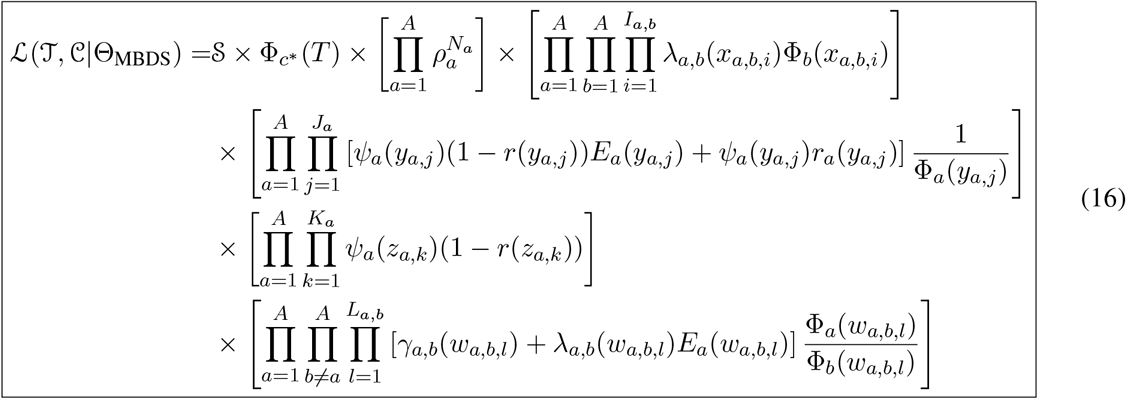

Here 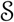 is an arbitrary form of conditioning as in equation (13) and Φ_*a*_(*τ*) = Ψ_*a*_(*τ,* 0), a complete list of notation is given in Table S4.

While the full colouring of the tree 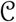 is never known, when fitting models where the diversification parameters are a function of a measurable state of the sampled nodes, the colour at some of the tip nodes, which we denote by 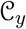, and at some of the internal ancestral samples, 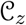, may be known. The likelihood of the tree topology 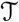 and these observed states can be calculated from equation (16) by integration.

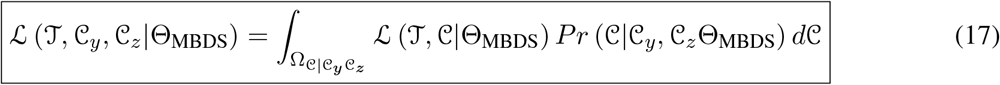

where 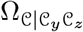 represents the space of all tree colourings that are consistent with the observed states. While it mathematically challenging to formulate the expression for 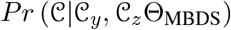 and hence this integral can not be done numerically it can be approximated via Monte Carlo integration. As exemplified for the BiSSEness model (Magnuson-Ford and Otto, 2012; Goldberg and Igić, 2012) in the supplementary material, one can simulate colourings within 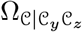 using stochastic mapping (Nielsen, 2002) proportional to their respective probabilities. Averaging the likelihood given by Equation (16) across these sampled coloured trees approximates the likelihood of a tree where only the state of sampled nodes are known.

Equations (16) and (17) provide a flexible definition of the tree likelihood, not only in terms of the diversification process but with respect to the specification of lineage states. As exemplified by the work of Barido-Sottani et al. (2020), the likelihood 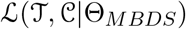 can be used to calculate model likelihoods where the state at the tips of the tree need not be specified a priori.

## Conclusion

Here we provide a unified framework for deriving birth-death-sampling models within both single- and mutli-type contexts. We derive a general likelihood expression for each of these cases, making as few assumptions about the processes that generated the data as possible. While drawing inferences from data will require making additional assumptions and applying mathematical constraints to the parameters (but see (Louca and Doebeli, 2018)), this framework can clarify the connections between various model variants, ease the development of new model variants tailored specifically to each situation, and provide a structure for understanding how results depend on model assumptions (Kirkpatrick et al., 2002; Lafferty et al., 2015; Louca and Pennell, 2020a). From a methodological perspective, our technique for deriving the likelihood of BDS models substantially lowers the barrier for developing and exploring new types of models in a way that Maddison *et al.* (Maddison et al., 2007) did for birth-death models without heterogeneous sampling. As evidence of this, the fully general and yet numerically tractable likelihood formula presented here allowed us to implement computational methods for simulating and fitting BDS models with arbitrary functional forms for *λ*, *μ*, *Ψ* and *r* (Louca and Doebeli, 2018). And importantly, given the recent discovery of widespread non-identifiability in birth-death processes fit to extant-only (Louca and Pennell, 2020a) and serially-sampled (Louca et al., 2021) phylogenetic data, there is a critical need to explore a much broader range of BDS models than were previously considered and the mathematical generalization presented in our paper will be enable this.

### Box 1 The connection between BDS and SIR models

The general single-type BDS model is intimately related to the SIR compartmental model used in classic theoretical epidemiology. This connection illustrates the explicit and implicit assumptions of the general BDS model and its sub models. Here we define the SIR epidemiological model, discuss how it can inform and be informed by these diversification models, and examine the shared assumptions of the two frameworks.

#### The SIR model

The SIR model is host-centric, partitioning the population via infection status into susceptible (S), infected (I), and recovered (R) hosts. Infection of susceptible hosts occurs at a per-capita rate *βI*. Infected hosts may recover (at rate *γ*), die of virulent cases (at rate *α*), or be sampled (at rate *Ψ*). The cumulative number of sampled hosts is represented in the SIR model shown in Figure B1 by *I*\*. Upon sampling, infected hosts may be treated and hence effectively recover with probability *r*. Hosts that have recovered from infection exhibit temporary immunity to future infection which wanes at rate *σ*. The special case of the SIR model with no immunity (the SIS model) is obtained in the limit as *σ* → ∞. In addition to these epidemiological processes, the SIR model includes demographic processes, such as host birth (rate *B*) and death from natural causes (rate *δ*). While not shown explicitly in the figure, these epidemiological and demographic rates may change over time as a result of host behavioural change, pharmaceutical and non-pharmaceutical interventions, or host/pathogen evolution.

#### The BDS Model

The BDS model is pathogen-centric, following the number of sampled and unsampled viral lineages over time, analogous to the *I* and *I*\* classes of the SIR model. A key element of general BDS model is that birth and death rates may vary over time. This time dependence may be either continuous (Morlon et al., 2011; Rabosky and Lovette, 2008b) or discrete (Stadler, 2011; Stadler and Bonhoeffer, 2013; Gavryushkina et al., 2014; Kühnert et al., 2014) Although arbitrarily time-dependent, the birth, death, and sampling rates in the general BDS model are assumed to be diversity-independent, analogous to the assumption of density-dependent transmission (pseudo mass action) in the SIR model (Keeling and Rohani, 2008). While some forms of diversity-dependence in diversification rates may be incorporated implicitly (Rabosky and Lovette, 2008a), explicit diversity-dependence (Etienne and Rosindell, 2012) goes beyond the scope of the BDS models considered here.

The general BDS model assumes all viral lineages are exchangeable - this has several implications. First, only a single pathogen type exists. Multi-type models (Stadler and Bonhoeffer, 2013; Kühnert et al., 2016; Barido-Sottani et al., 2020, 2018) are not included in the GBDS framework. Second, transmission is independent of lineage age. In the macroevolutionary case, such age-dependence has been suggested to reflect niche differentiation in novel species (Hagen et al., 2015) and in the epidemiological case may reflect adaptation towards increased transmissibility following a host species-jumping event. Third, lineage exchangeablity is reflected in the absence of an exposed (E) class in the SIR model in which hosts can, for example, transmit infections but not be sampled or vice versa. Finally, the general BDS model assumes all lineages are sampled at random and does not include sub-models with non-random representation of lineages (Stadler et al., 2012).

**Figure B1:**
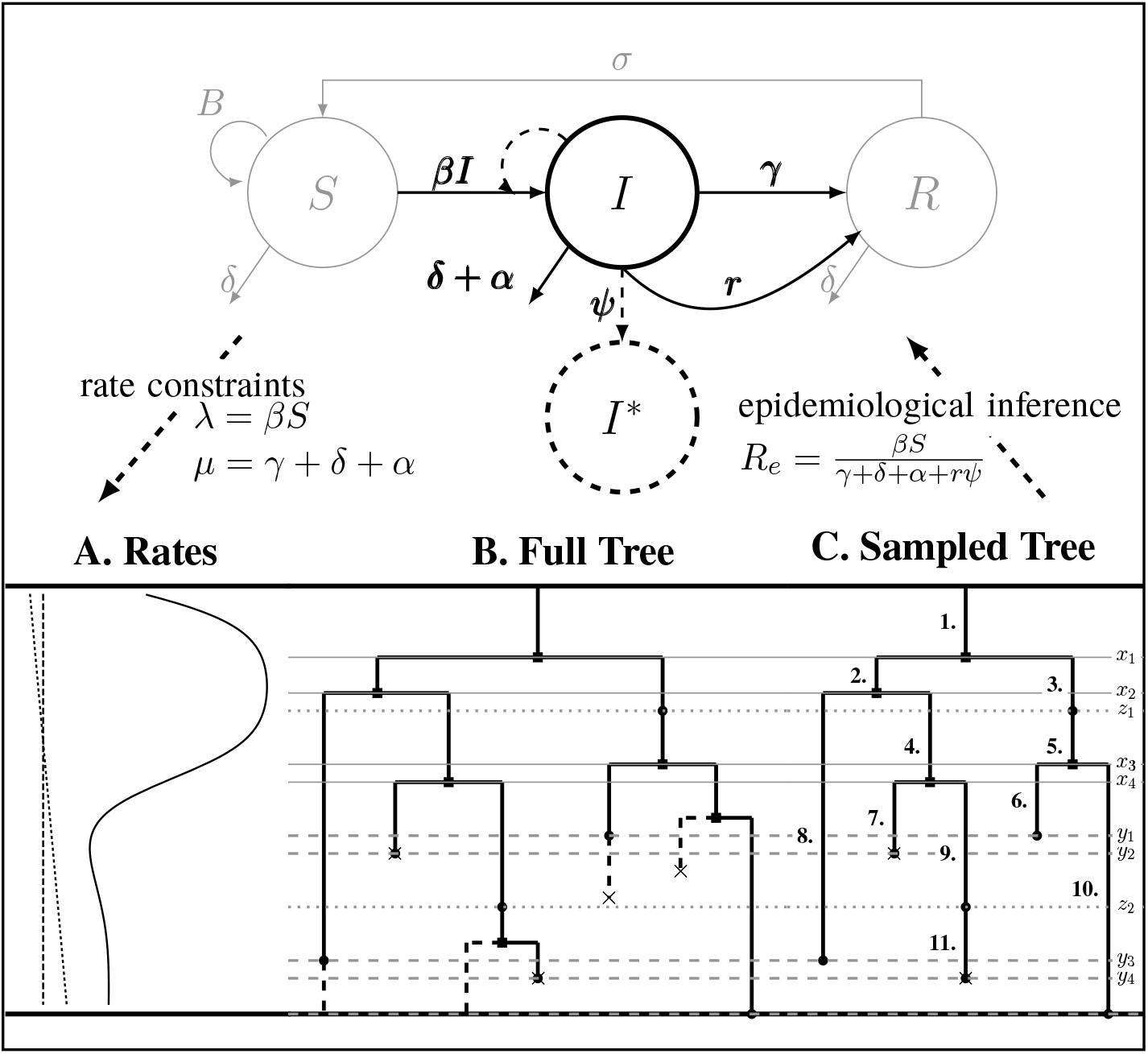
Top: The SIR epidemiological model. Black (gray) lines and classes represent rates and variables followed (in)directly by the BDS model. The SIR model can be used to constrain the rates of the BDS model (A.). Simulated forward in time, the result of the BDS stochastic processes is a *full tree* (B.) giving the complete genealogy of the viral population. Pruning away extinct and unsampled lineages produces the *sampled tree* (C.). Arising from a BDS process, this sampled tree can be summarized in two ways. First by the set of edges (labeled 1-11) or as a set of critical times (horizontal lines) including: 1) the time of birth events (solid, *x*_*i*_) 2) terminal sampling times (dashed, *y*_*j*_), and 3) ancestral sampling times (dotted, *z*_*k*_). Given the inferred rates from a reconstructed sampled tree, these rates can be used to estimate characteristic parameters of the SIR model, for example the basic or effective reproductive number.

## Model Connections

Given their shared model assumptions, the general BDS model can be constrained explicitly to reflect an underlying SIR epidemic by setting the viral birth rate equal to the per-capita transmission rate of the infectious class *λ*(*τ*) = *βS*(*τ*) and the viral death rate to the infectious recovery or removal rate *μ*(*τ*) = *γ* + *δ* + *α*, whereas the sampling rate *Ψ*(*τ*) is identical across models. While constraining the birth, death, and sampling rates in this manner can be used to parameterize compartmental models (Kühnert et al., 2014) doing so is an approximation assuming independence between the exact timing of transmission, recovery or removal from population, and sampling events in the SIR model and birth, death, and sampling events in the diversification model. The resulting tree likelihood in terms of the compartmental model is given by:

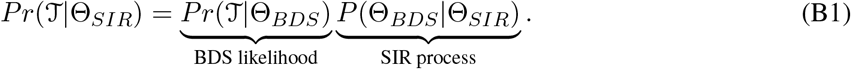

While they are not sub-models of the general BDS process, likelihood models have been developed that capture the full non-independence of viral diversification and epidemiological dynamics for the SIR model specifically (Leventhal et al., 2012) and in compartmental models in general (Vaughan et al., 2019). The connection between the BDS process and SIR epidemiological models can also be used after the diversification rates are inferred to estimate the basic and effective reproductive rates (Stadler et al., 2012; Stadler and Bonhoeffer, 2013). Specifically, the effective reproductive rate at time *τ* before the present day is given by 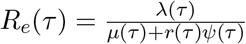

## Acknowledgments

We would like to thank Sally Otto for her thoughtful comments on this work. This work was supported by a *Grant for Catalyzing Research Clusters* awarded to the UBC Biodiversity Research Centre, NSF DEB Grant #2028986 award to SL and MWP. MWP was supported by a NSERC Discovery Grant and AM was supported in part by the EEB department Postdoctoral Fellowship from the University of Toronto. AMcL was supported in part by a Canadian Institutes of Health Research (CIHR) doctoral award (#6557). JBJ is supported by a Genome Canada Bioinformatics and Computational Biology grant (287PHY), CIHR coronavirus rapid response program grant (440371) and is grateful to the British Columbia Centre for Excellence in HIV/AIDS for additional funding support.

## Appendix: Adding assumptions to the general model

In this appendix, we demonstrate how one can obtain the likelihood of sub-models with different sets of assumptions by applying constraints to the general likelihood. There are four classes of assumptions that are commonly applied in epidemiological and macroevolutionary studies. First, researchers can make assumptions about the functional form of the birth, death, and sampling rates. Here we address two such unique assumptions: i) Birth, death, and sampling rates are constant (see section 1, S1.1, S1.2, S1.5); and ii) birth, death, and sampling rates are piecewise-constant functions of time (see sections 1, S1.6). The cases where birth, death, and sampling rates are defined by a stochastic or deterministic SIR model are mathematically analogous to the cases of the piecewise-constant and general time-variable models respectively. All additional constraints imposed will depend on the exact compartmental model used and hence we will not discuss them in detail in this section. The second major class of assumptions pertains to sampling. There are four such sampling assumptions: i) sampling happens only at the present day as in a birth-death model (see section 1, S1.1, S1.3, S1.4) or as implemented in the “Birth Death Skyline Contemporary” prior in the BDSKY package in BEAST; ii) the absence of concerted present-day sampling (see section 1, S1.5); iii) the inclusion of ancestral samples with sampled descendants (S1.7, and S1.6); and iv) concerted sampling attempts (CSA) during which all lineages are sampled with a given probability (see sections 1, S1.6). The third assumption class considers the presence of mass extinction events (see section 1, S1.5). The fourth and final major class of assumptions deal with the conditioning of the likelihood. The various conditioning schemes are explored in section 1 and summarized in Table S3.

### Rate assumptions

#### Constant rates

- *Model Assumptions:* Constant diversification rates: *λ*(*t*) = *λ*, *μ*(*t*) = *μ*, *Ψ*(*t*) = *Ψ*, and constant removal probability *r*(*t*) = *r*.
- *The IVP for g*_*e*_(*τ*):

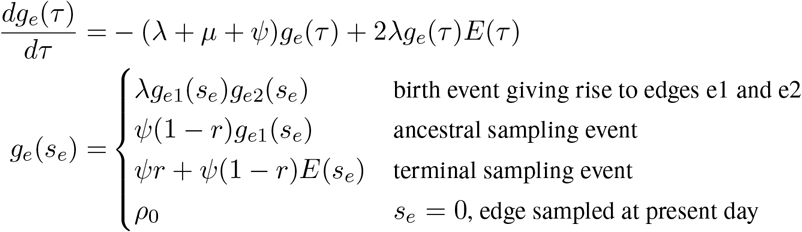
- *The IVP for E*(*τ*):

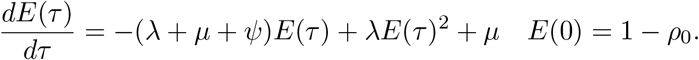 In this case the IVP for *E*(*τ*) is a Bernoulli differential equation and has a known analytical solution. As given by equation 1 in Stadler (2010) this solution is given by:

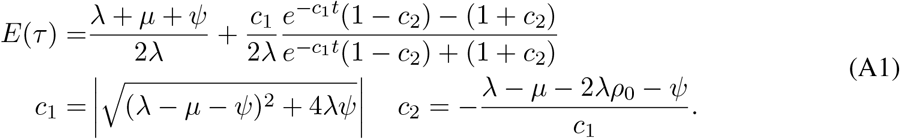
- *The Probability Flow:*

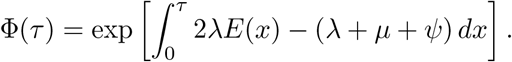
- *The Likelihood:*

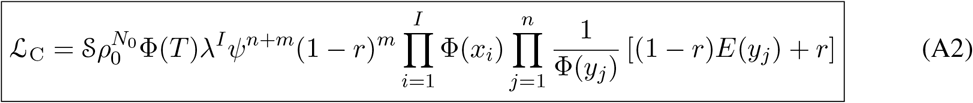

#### Piecewise-constant rates

- Model assumptions: Divide time into *L* + 1 intervals defined by *transition times* 0 = *t*_0_ *< t*_1_ *< t*_2_ *< ... < t*_*L*_ < *t*_*L*+1_ = *T*. Define rates and removal probabilities constant within a given interval.

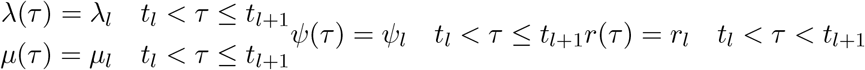
- *The IVP and Solution for g*_*e*_(*τ*): Given the definitions of *λ*(*τ*), *μ*(*τ*), *Ψ*(*τ*), and *r*(*τ*) above the IVP for *g*_*e*_(*τ*) is identical to that given in Equations (3) and (4).
- *The IVP and Solution for E*(*τ*): As with *g*_*e*_(*τ*), the IVP for *E*(*τ*) is given by Equation (8). With the piecewise-constant rate assumptions, however, the general solution for *E*(*τ*) between *t*_*l*_ < *τ* ≤ *t*_*l*+1_ is known (similar to Equation (A1)). Defining *E*_*l*_(*τ*) = *E*(*τ*) where *t*_*l*_ < *τ* ≤ *t*_*l*+1_ and *E*_*l*_(*t*_*l*_) = *E*_*l*−1_(*t*_*l*_) we have:

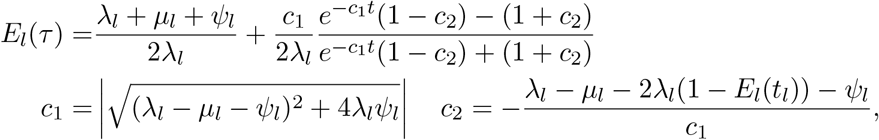

where *E*_*l*_(*t*_*l*_) = *E*_*l*−1_(*t*_*l*_) for *l >* 0 and *E*_0_(*t*_0_) = 1 − *ρ*_0_.
- *The Probability Flow:* We define a probability *sub-flow* within each time interval. Specifically, in the *l*^*th*^ time interval.

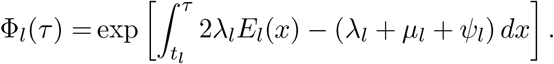 The complete flow can be expressed as a function of the sub-flows in the following manner:

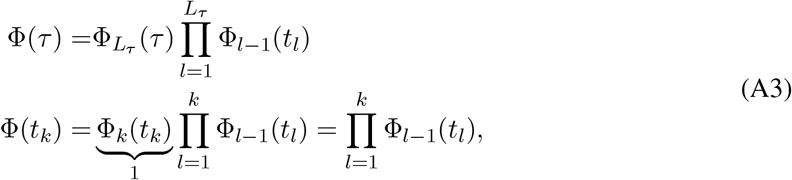

where *L*_*t*_ is the index of the time *t*_*l*_ at or after time *t*, i.e. the largest index such that *t*_*l*_ ≤ *τ*.
- *The Likelihood:* Given these piecewise definitions we substitute them into the general BDS likelihood (13).

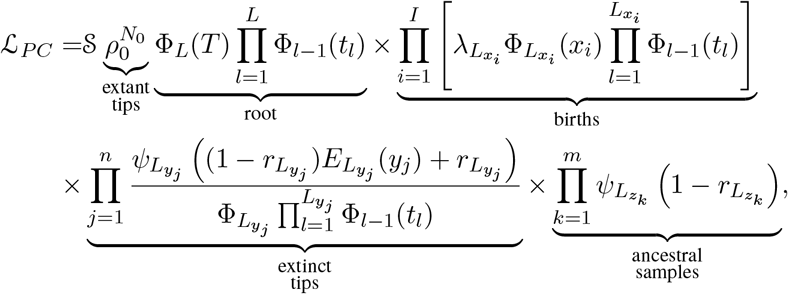

where we use *PC* to denote the piecewise-constant assumption. We can simplify several of these products. Let *α*_*l*_ be the number of birth events ≥ *t*_*l*_ and *σ*_*l*_ the number of sampling events ≥ *t*_*l*_.

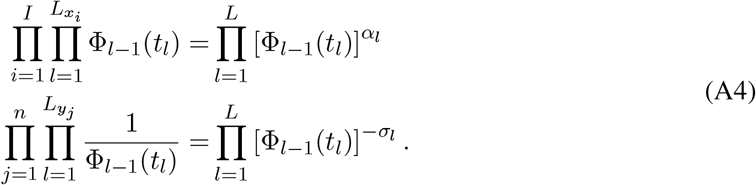 Let *n*_*l*_ be the number of observed lineages alive at time *t*_*l*_. Because the number of observed lineages increases with each birth and decreases with each sampled tip, counting the root we have *n*_*l*_ = *α*_*l*_ − *σ*_*l*_ + 1. Substituting the expressions for the into the likelihood and using the definition of *n*_*l*_ we have:

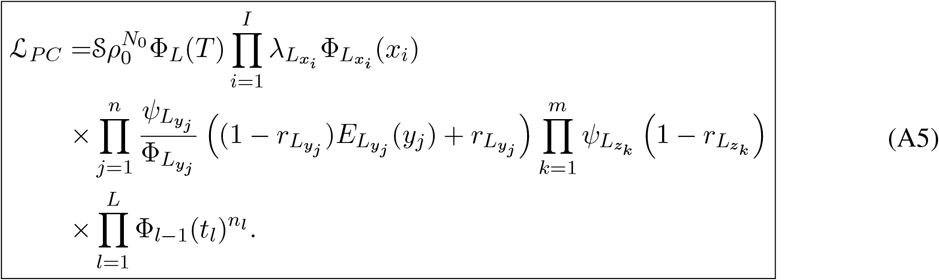

### Sampling assumptions

#### Birth-death models

- *Model Assumptions:* The birth-death model assumes that *Ψ*(*τ*) = 0. Note that the probability of sampling a lineage given it is alive at the present day remains as *ρ*_0_ (incomplete sampling).
- *IVP for g*_*e*_(*τ*):

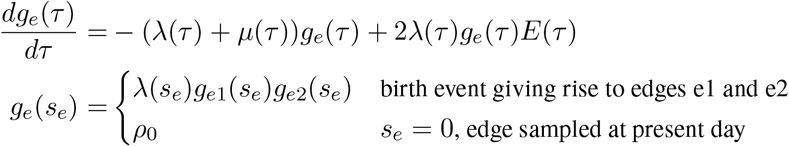
- *IVP for E*(*τ*):

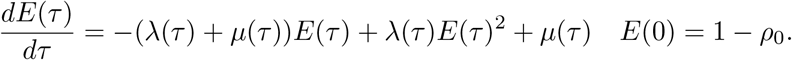 Note in this case *E*(*τ*) equals *Ê*(*τ*), the probability a lineage leaves no sampled extant descendants. As demonstrated by Morlon et al. (2011) there exists a general solution to this initial value problem, see section S1.3 for more details. This general solution is given by:

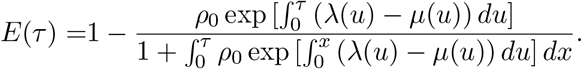
- *The Probability Flow:* From (Morlon et al., 2011), the probability flow can be written as the following:

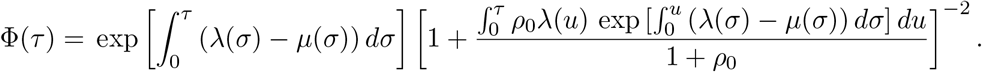
- *The Likelihood:*

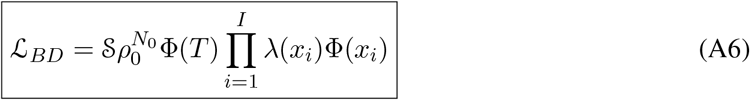

#### No sampling at the present

Here we consider the case when *ρ*_0_ = 0. The likelihood follows exactly as in the general model case. The resulting likelihood expression is given by:

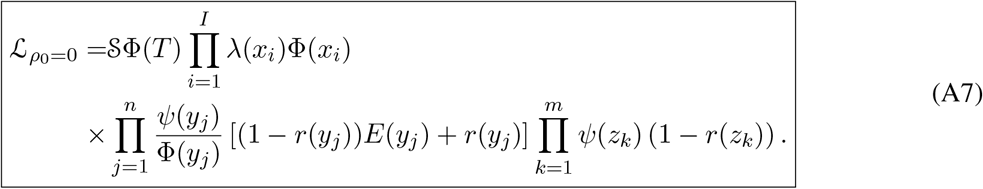

Note that in this case *I* = *n* − 1.

#### Concerted sampling attempts

- *Model Assumptions:* Here we introduce *L* concerted sampling attempts (CSA) at known points in time, *t*_*l*_ *l* ∈ {1, 2*, ...L*}. Like the CSA at the present day, and in contrast to the background Poissonian sampling rate, during the CSA at time *t*_*l*_ every lineages is sampled with a fixed probability *ρ*_*l*_. In the derivation of the likelihood below, we must distinguish between three different sampling event types. First, *past Poissonian sampling events* are those that do not occur during CSAs. Second, *past concerted sampling events* are those that occur during a CSA at time *t*_*l*_ *l* ∈ {1, 2*, ..., L*}. Finally, *present concerted sampling events* are those that occur at the present day *τ* = 0. Past concerted sampling attempts can be included in the general model above by adding *L* Dirac distributions to the Poisson sampling rate function. Namely,

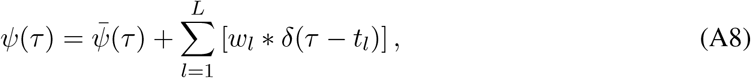

where 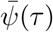 is the background Poissonian sampling rate and *w*_*l*_ = −*ln* (1 − *ρ*_*l*_). The definition of *w*_*l*_ comes from solving the CDF of the exponentially distribution for the ‘effective sampling rate’ such that the probability of a lineage being sampled is *ρ*_*l*_.
- *IVP for g*_*e*_(*τ*):

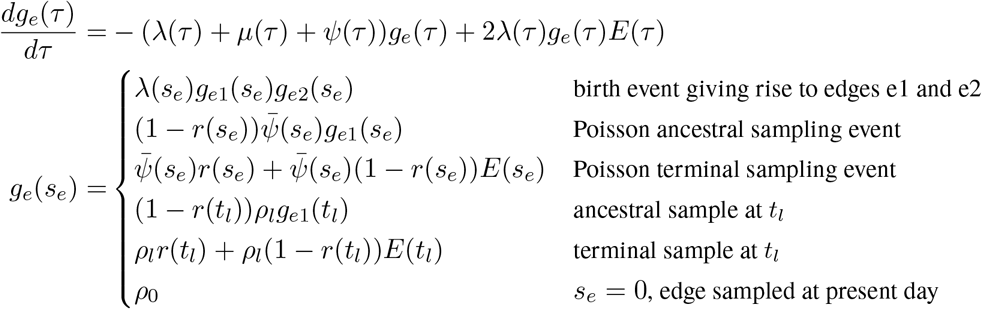 The solution to *g_e_*(*τ*) is given by Equations (5) and (6).
- *IVP for E*(*τ*): As with *g*_*e*_(*τ*), the IVP for *E*(*τ*) is identical to that given for the general model in Equation (8). Except in rare cases the IVP must be solved numerically hence requiring numerical integration over Dirac distributions which can prove to be problematic. Note however, that when examining the integrals over the CSAs, a priori, it is a matter of convention whether the Dirac distribution should be considered as “integrated over” when located at the upper integration bound 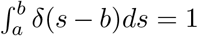 or at the lower integration bound 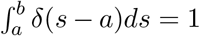. Whichever convention we chose, we must rigorously obey it so that the ratio Φ(*t*)/Φ(*s*) correctly evaluates to Ψ(*s, t*) whenever *s ≤ t*. Using the former convention, we can rewrite the probability *E*(*t*_*l*_) at each concerted sampling time *t*_*l*_ as:

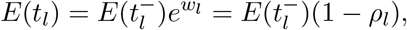

where 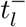 denotes the limit as time approaches *t*_*l*_ from below. Hence the probability *E*(*τ*) at any time *τ* can be evaluated numerically by considering the dynamics between successive CSAs and at each CSA separately.
- *The Probability Flow:* The probability flow is given by:

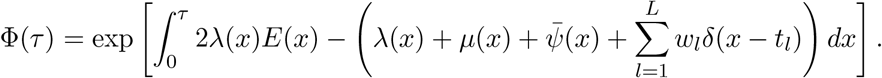 As with *E*(*τ*) integration over the dirac distributions can be problematic and hence we rewrite this expression separating out these terms. Let *L*_*τ*_ be the oldest CSA occurring at or after time *τ*, i.e. the largest index for which *t*_*l*_ ≤ *τ*.

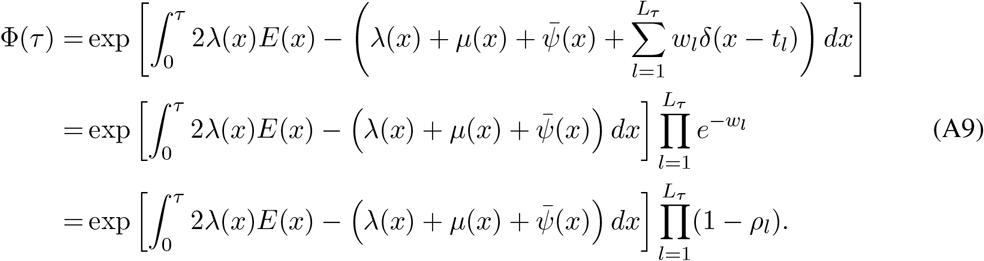 We define:

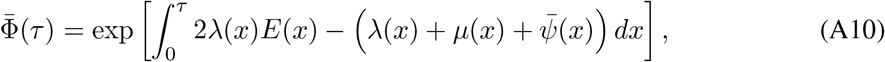

which means that we can rewrite Equation (A9) as:

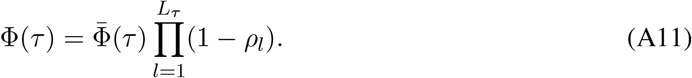
- *The Likelihood:* The edge representation of *g*_*stem*_ is given by:

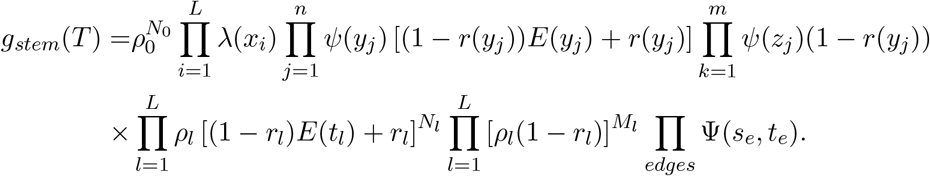 The critical time representation of *g*_*stem*_ is given by:

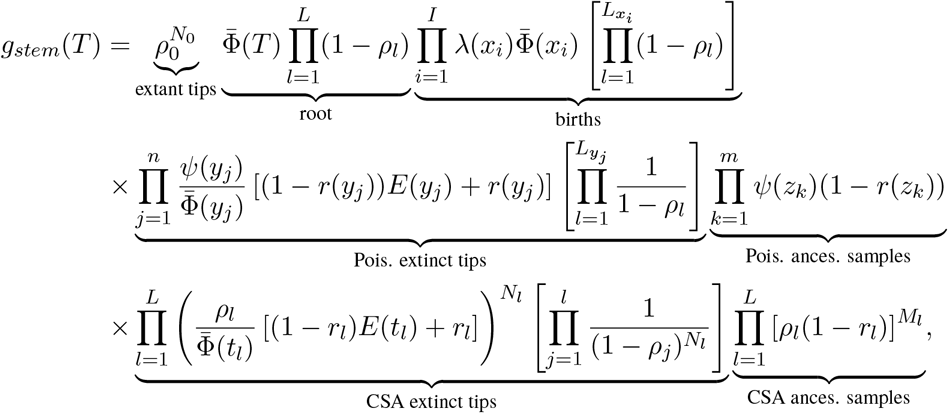

where *N*_*l*_ is the number of tip samples (samples without descendants) obtained during the *l*^*th*^ CSA and *M*_*l*_ is the number of ancestral samples (sequences with descendants). By changing how we enumerate birth, death, and sampling events we can greatly simplify this likelihood. First, let *α*_*l*_ be the number of branching events at or before the the *l*^*th*^ CSA. In other words, *α*_*l*_ is the number of branching events if the tree were trimmed at the *l*^*th*^ CSA. Then:

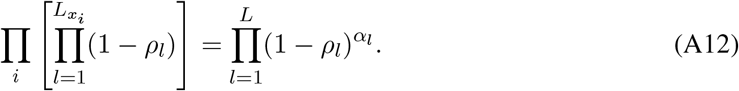 Second, let *σ*_*l*_ be the number of past Poissonian sampling events before time *t*_*l*_. Then:

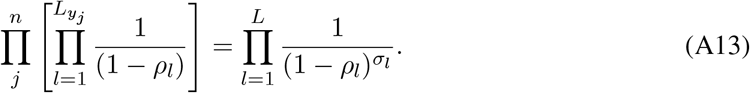 Finally, let *β*_*l*_ be the number of past lineages sampled during a CSA at or before the CSA at time *t*_*l*_. Hence, *β*_*l*_ = *N*_*l*_ + *N*_*l*+1_ + ... + *N*_*L*_. Then:

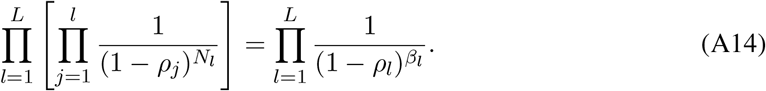 The likelihood hence simplifies to:

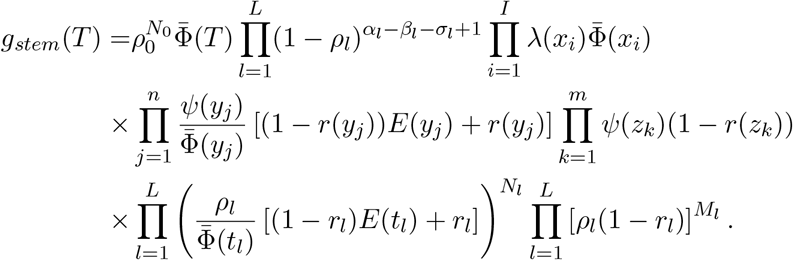 Let *n*_*l*_ be the number of lineages that cross *t*_*l*_, i.e., the number of lineages alive at time *t*_*l*_ with sampled descendants at some younger age. Note that by this definition *n*_0_ = 0. Then *b*_*l*_ + *σ*_*l*_ + *n*_*l*_ is the number of tips in the tree had it been trimmed at age *t*_*l*_ whereas *α*_*l*_ is the number of branching events. Therefore we must have *α*_*l*_ = *b*_*l*_ + *σ*_*l*_ + *n*_*l*_ − 1. This allows us to simplify the conditioned likelihood given below.

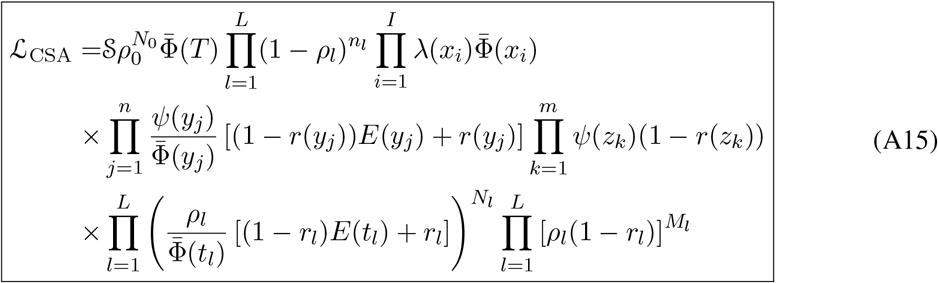

### Mass extinction

- *Model Assumptions:* In addition to the Poisson birth death and sampling events considered in the general model, there are *L* mass extinctions events occurring at times *t*_1_ *> t*_2_ *> ..t*_*L*_. During the *l*_*th*_ mass extinction event each lineage goes extinct with probability *ν*_*l*_. As with concerted sampling such mass extinction events can be introduced into the model by adding a set of dirac-delta functions to the Poisson death rate, 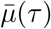.

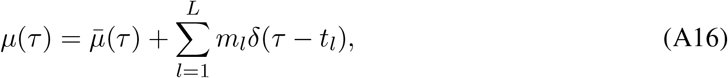

where *m*_*l*_ = −*ln*(1 − *ν*_*l*_).
- *IVP for g*_*e*_(*τ*): The initial value problem for *g*_*e*_(*τ*) is identical to that given in equation by Equations (3) and (4) except that *μ* is now includes the mass extinction events.
- *IVP for E*(*τ*): The IVP for *E*(*τ*) to that given by Equation (8) except where the extinction rate is given by Equation (A16). The solution to *E*(*τ*) is obtained by numerical integration. Given the dirac-delta functions this numerical integration can be carried out in a piecewise manner integrating separately between and over each mass extinction event. Defining 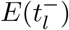 as the solution up to but not including the mass extinction event at time *t*_*l*_ we have:

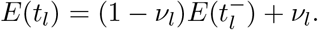 The first term reflects the probability that a lineage that does not go extinct during the *l*^*th*^ mass extinction event leaves no observable offspring (with probability 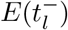) whereas the second term reflects the fact that all lineages that go extinct during the *l^th^* mass extinction leave no observed descendants with probability 1.
- *The Probability Flow:* The solution to the IVP is once again given by 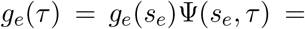 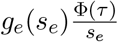 where:

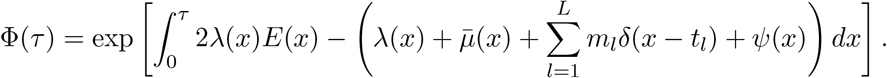 As with the CSAs, let *L*_*τ*_ be the last index *l* such that *t*_*l*_ < *τ*. We can separate out the mass extinction terms in the following way.

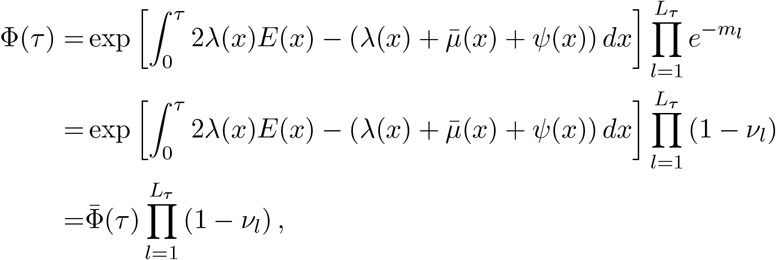

where 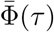 is defined as in Equation (A10).
- *The Likelihood:* Given these initial value problems the likelihood follows as in the general model.

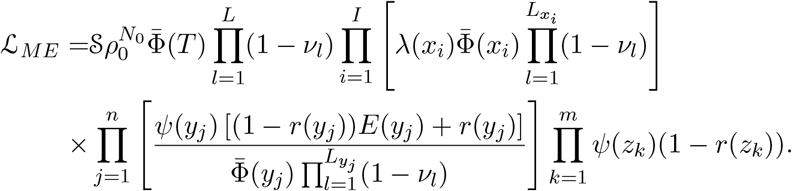 As with the CSAs we can use relations analogous to Equations A12–A14 to rewrite the likelihood:

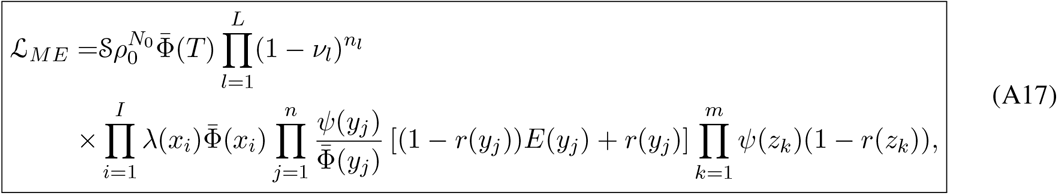

where *n*_*l*_ is defined as before as the number of lineages present at time *t*_*l*_.

### Alternative conditioning

Table S3 lists a number of possible conditionings, 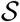 that can be applied to the tree likelihood. First, is the trivial case of no conditioning 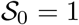 which gives the likelihood of the observed tree including the stem edge between time *T* and *t*_MRCA_. The likelihood of the phylogeny excluding the stem edge, i.e., conditioning of the *t*_MRCA_, can be obtained by setting 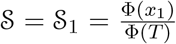. Recall that the elements of 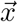 are ordered such that *x*_1_ = *t*_MRCA_ is the first (oldest) birth event.

Acknowledging that one would not reconstruct a phylogeny without any sampled lineages, we can condition the likelihood on observing at least one sampled lineage (either at or before the present day) given the time of origin, 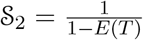. Or as with 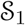, conditioning on at least one sampled lineage given the *t*_MRCA_. In order to have at least one sampled lineage *and* a most recent common ancestor, however, each daughter lineage of the common ancestor must have at least one descendent. Hence we have 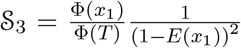. The general birth-death-sampling model assumes that all lineages alive at the present day are sampled with probability *ρ*_0_. As with concerted sampling attempts (CSAs) prior to the present day, this present day CSA may include the sampling of multiple lineages as well as possibly resulting in no sampled lineages. As with 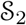 and 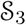 we can condition the tree likelihood on observing at least one extant lineage at the present day. To do so, we define *Ê*(*τ*) = *E*(*τ|Ψ* = 0), the probability that a lineage alive at time *τ* has no extant descendants. Conditioning on the time of origin we have: 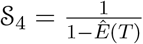. Conditioning on the time of the most recent common ancestor we have: 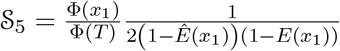, where now at least one of the two daughter lineages of the common ancestor has a present day sample. In many cases 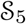 is modified, however, to condition on *both* daughter lineages having an extant sampled descendent: 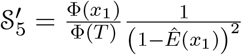.

As an alternative to conditioning on at least one extant sampled descent, tree likelihoods can be conditioned on having *exactly N*_0_ sampled (extant) descendants. Let 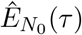 be the probability a lineage alive at time *τ* has exactly *N*_0_ descendants. Although a general expression for 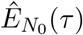 is unknown, in the case of constant birth, death, and sampling rates (the case in which this form of conditioning has been applied), the expression for 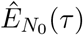 is given by (Gernhard, 2008; Kendall, 1948) and **Theorem 3.3** by Stadler Stadler (2010):

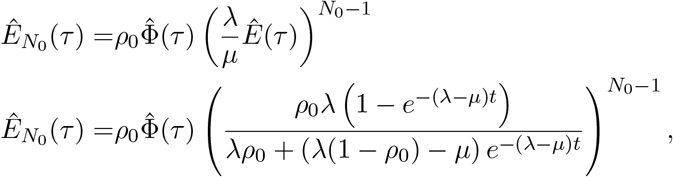

where, like *Ê*, 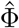 is given by Equation (10) evaluated with where *Ψ* = 0. Given the time of origin we can condition on observing exactly *N*_0_ lineages by setting 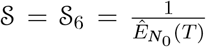. When *t*_MRCA_ is given instead, then the number of descendants of the two daughter lineages must add up to *N*_0_ while both daughter lineages must still have at least one descendant(see Stadler (2010) **Corollary 3.9**).

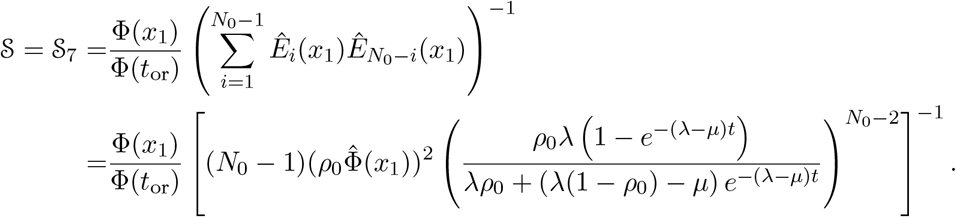

While early BDS models often employed such conditioning (Stadler, 2009, 2010), this form of conditioning has not been employed in many later models perhaps because the biological justification for such conditioning is vague.

The final form of conditioning used in the literature, which we will represent simply as 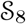, is the multiplication of the BDS likelihood by a constant to account for the enumeration over the possible indistinguishable representations of a given tree. The value of this constant depends on whether the tree considered is “labeled” and “oriented” (Gavryushkina et al., 2013) and whether, as in the derivation here, the vector of birth events, 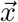, is (un)ordered. Inclusion of such a constant should have no effect on the maximum likelihood inference of the model parameters given a specified tree. In cases where the constant is a function of the critical times (Heath et al., 2014), it can influence the inference when the parameters and the tree are jointly estimated.

## Part I

### Single-type model

#### S1 Relationships between existing models

In the previous section, we proved that one could go from the most general case to specific sub-models by incorporating additional constraints to the parameters. In this section, we illustrate how to work in the other direction — that is, to start with the most assumptions of a particular sub-model and derive its likelihood function using the same five-step procedure used to derive the general BDS model in the Main Text. In addition to illustrating the utility of our mathematical technique, by deriving the likelihoods of previously developed models, we are able to unify a diverse and, occasionally opaque, literature using a common terminology, notation, and formulation.

##### S1.1 Stadler 2009

Here we re-derive the likelihood given by **Equation 2** in (Stadler, 2009). Note throughout all equation, corollary, and theorem references in other publications will be placed in bold face type.

- Step 1: Specify the model.

– Constant rates: *λ*(*τ*) = *λ*, *μ*(*τ*) = *μ*.
– Birth-death model with incomplete sampling at present day: *Ψ*(*τ*) = 0 and *ρ*_0_ ≤ 1.
– Conditioning on there being exactly *N*_0_ lineages at the present day given the time of origin, 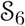 and un-ordered birth events 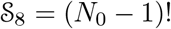.
- Step 2: IVP for *g*_*e*_(*τ*).

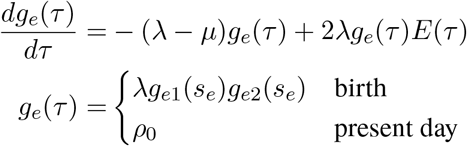
- Step 3: IVP for *E*(*τ*).

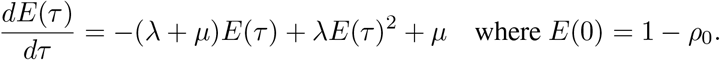 Given the constant rate assumption there exists a general solution for *E*(*τ*).

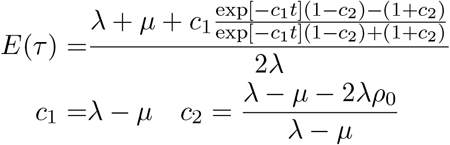
- Step 4: Derive *g*_*stem*_(*T*).

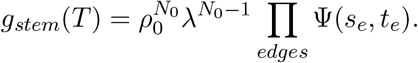
- Step 5: Calculate *g*_*stem*_(*T*) wrt the critical time representation. Given the assumption of constant rates and no Poisson sampling the expression for Φ(*τ*) simplifies to:

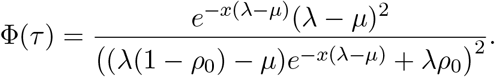 Hence we have:

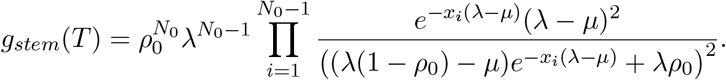
- Step 6: Impose conditioning. The likelihood is conditioned on having exactly *N*_0_ lineages at the present day 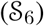 and a constant 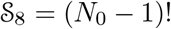 as the birth events are left un-ordered.

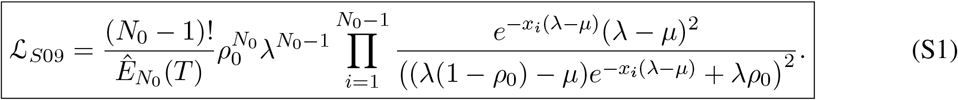

##### S1.2 Stadler 2010

- Step 1: Specify the model.

– Constant rates: *λ*(*τ*) = *λ*, *μ*(*τ*) = *μ*, *Ψ*(*τ*) = *Ψ*.
– No removal upon sampling, *r* = 0.
– Multiple presented.

– **Equation 3** (Stadler, 2010): No conditioning.
– **Equation 4** (Stadler, 2010): Exactly *N*_0_ extant sampled tips.
– **Corollary 3.7** (Stadler, 2010): At least one extant tip conditioning on the time of origin.
– **Equation 5** (Stadler, 2010): At least one extant tip conditioning on the *t*_*MRCA*_.
– **Equation 6** (Stadler, 2010): *N*_0_ extant tips conditioning on the time the *t*_*MRCA*_.
- Step 2: IVP for *g*_*e*_(*τ*).

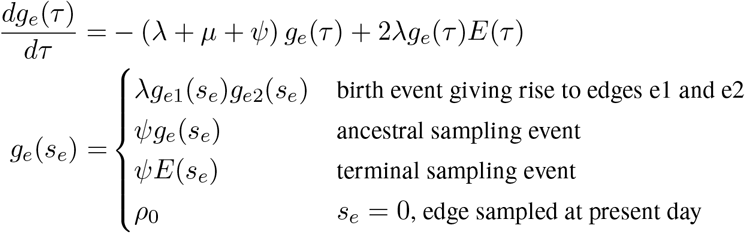
- Step 3: IVP for *E*(*τ*). Given the constant rate assumption there exists a general solution for *E*(*τ*).

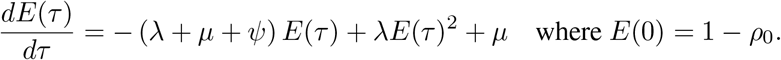 This is a Bernoulli differential equation and has a known solution as given by Stadler (Stadler, 2010).

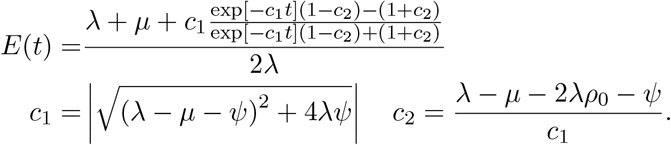
- Step 4: Derive *g*_*stem*_(*T*).

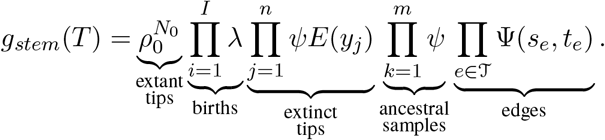
- Step 5: Calculate *g*_*stem*_(*T*) wrt the critical time representation. Given the assumption of constant rates and no Poisson sampling the expression for Φ(*τ*) simplifies to:

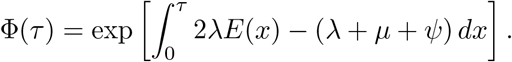 Hence we have:

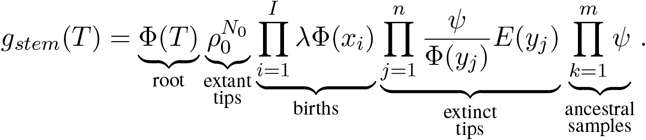
- Step 6: Impose conditioning. Imposing an arbitrary conditioning we have the following likelihood. Note that *I* = *N*_0_ + *n* − 1.

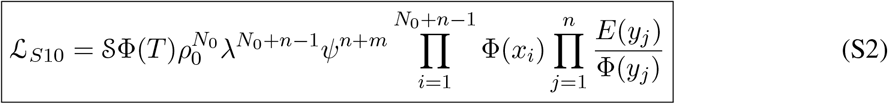

– **Equation 3** (Stadler, 2010): No conditioning. 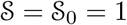.
– **Equation 4** (Stadler, 2010): Exactly *N*_0_ extant sampled tips. 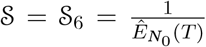 where in the case of constant rates we have:

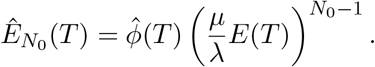

where 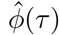 equals *φ*(*τ*) where *Ψ* = 0.
– **Corollary 3.7** (Stadler, 2010): At least one extant tip conditioning on the time of origin. 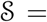 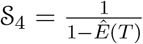 where *Ê*(*T*) is given by the solution for *E*(*τ*) above given that *Ψ* = 0.
– **Equation 5** (Stadler, 2010): At least one extant tip conditioning on the *t*_*MRCA*_, 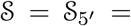 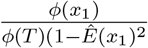.
– **Equation 6** (Stadler, 2010): *N*_0_ extant tips conditioning on the time the *t*_MRCA_, 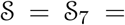 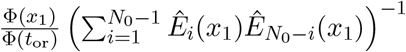,

##### S1.3 Morlon *et al.* 2011

Here we derive **Equation 1** from (Morlon et al., 2011).

- Step 1: Specify the model.

– Time variable rates.
– Birth-death only *Ψ*(*τ*) = 0.
– At least one extant sample 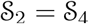.
- Step 2: IVP for *g*_*e*_(*τ*).

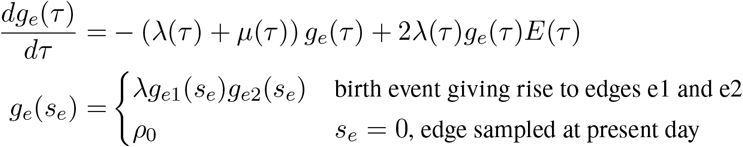
- Step 3: IVP for *E*(*τ*).

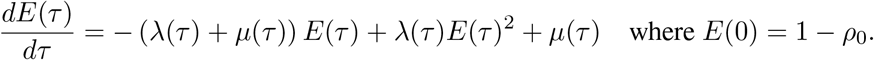 The general solution of this differential equation is given by:

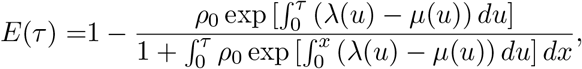

see **Equation 2** in (Morlon et al., 2011).
- Step 4: Derive *g*_*stem*_(*T*).

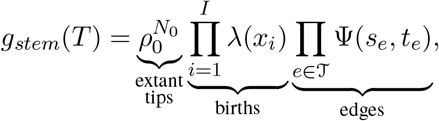

where the expression for Ψ(*s*_*e*_, *t*_*e*_) is given by **Equation 3** in (Morlon et al., 2011).
- Step 5: Calculate *g*_*stem*_(*T*) wrt the critical time representation. Given Φ(*τ*) = Ψ(0*, τ*) from **Equation 3** for in (Morlon et al., 2011) we have:

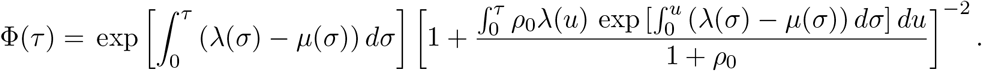 Hence we have:

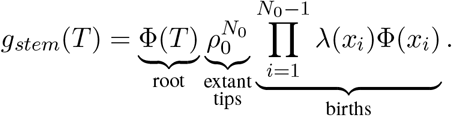 Note *I* = *N*_0_ − 1.
- Step 6: Impose conditioning. The likelihood given by **Equation 1** in (Morlon et al., 2011) is conditioned on the existence of at least one sampled lineage, 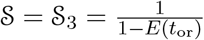.

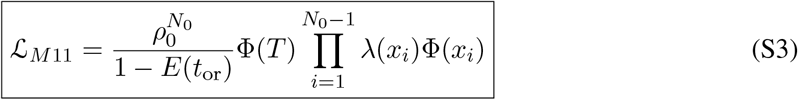

##### S1.4 Stadler *et al.* 2011

Here we derive the likelihoods given by **Theorem 2.6** and **2.7** in (Stadler, 2011).

- Step 1: Specify the model.

– piecewise-constant Poissonian rates. *λ*(*τ*) = *λ*_*l*_ and 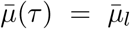 if *t*_*l*_ ≤ *τ* < *t*_*l*+1_ *l* = 0, 2, ...*L* + 1 where *t*_0_ = 0 and *t*_*L*+1_ = *T*.
– No Poisson sampling, *Ψ*(*τ*) = 0.
– Mass extinction events at times *t*_*l*_ *l* = 1, 2*, ...L* as specified above.

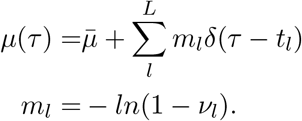
– **Theorem 2.6** (Stadler, 2011) imposes no additional conditioning 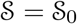 whereas **Theorem 2.7** (Stadler, 2011) conditions on observing at least one descendent given the time of the most recent common ancestor 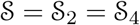.
- Step 2: IVP for *g*_*e*_(*τ*).

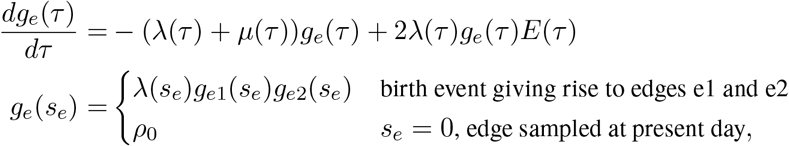

where *μ*(*τ*) is given above. In terms of the probability flow *g*_*e*_(*τ*) = *g*_*e*_(*s*_*e*_)Ψ(*s*_*e*_, *τ*), where:

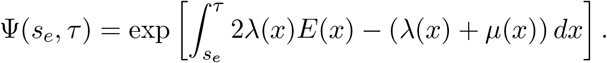 Here *μ*(*τ*) includes the mass extinction events.
- Step 3: IVP for *E*(*τ*).

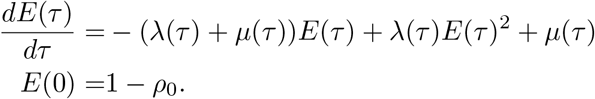 Given the piecewise constant nature there is a known general solution. Let *E*(*τ*) = *E*_*l*_(*τ*) where *t*_*l*_ < *τ* ≤ *t*_*l*+1_. Then define 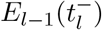 as the solution up to but not including the mass extinction event at time *t*_*l*_ we have:

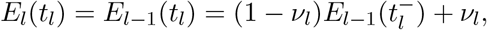

where *E*_*l*_(*τ*) is given by a solution similar to that in Equation (A1).

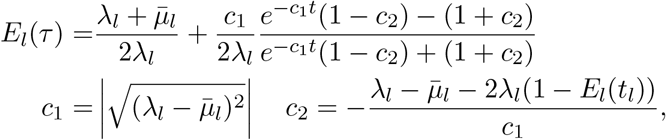

where *E*_0_(*t*_0_) = 1 − *ρ*_0_.
- Step 4: Derive *g*_*stem*_(*T*). The expression for *g*_*stem*_(*T*) is given by:

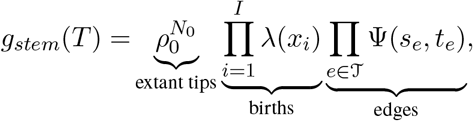

where *λ*(*τ*) and Ψ(*s*_*e*_, *τ*) are specified above.
- Step 5: The critical time representation. We once again define the sub-flow Φ_*l*_(*τ*) where *t*_*l*_ < *τ* ≤ *t*_*l*+1_:

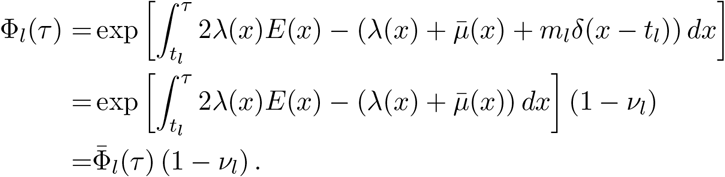 Note *ν*_0_ = 0. The complete flow is, as given by Equation (A3).

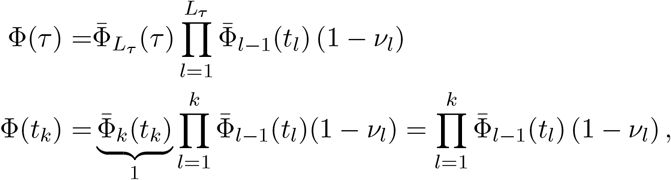

where *L*_*τ*_ is once again the largest index *l* such that *t*_*l*_ < *τ*. The critical time representation of *g*_*stem*_ then is:

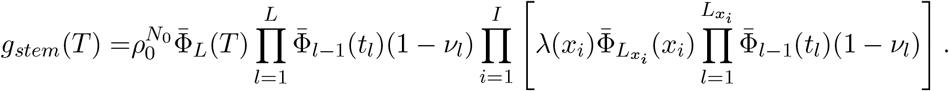 Defining *α*_*l*_ as the number of observed birth events ≥ *t*_*l*_ we can rewrite the product:

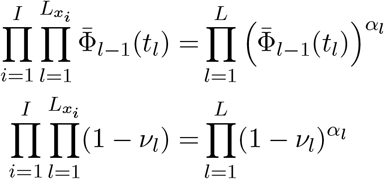 Hence we have:

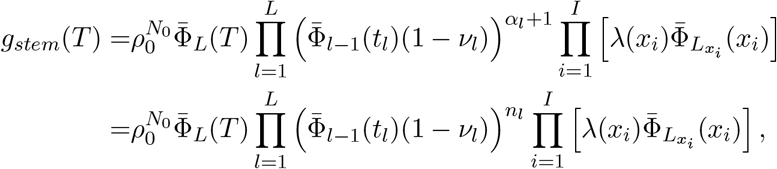

where *n*_*l*_ is the number of lineages in the observed phylogeny at time *t*_*l*_ then *n*_*l*_ = *α*_*l*_ + 1.
- Step 6: Likelihood conditioning. For **Theorem 2.6** (Stadler, 2011) the likelihood is given by:

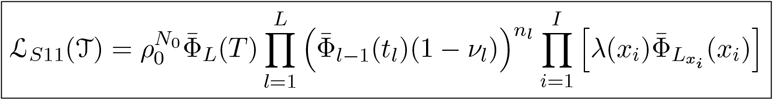 For **Theorem 2.7** (Stadler, 2011) the likelihood is given by:

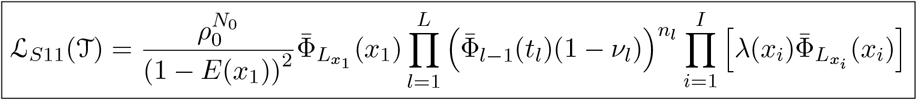

##### S1.5 Stadler *et al.* 2012

Here we derive **Equation 1** in (Stadler et al., 2012).

- Step 1: Specify the model.

– Constant birth, death, and sampling rates.
– No present day sampling. All lineages removed upon sampling *r* = 1.
– No conditioning.
- Step 2: IVP for *g*_*e*_(*τ*).

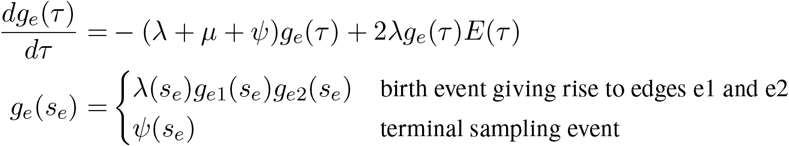
- Step 3: IVP for *E*(*τ*).

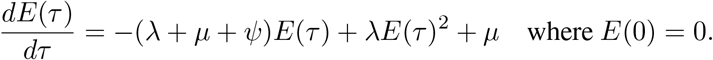 The solution to this differential equation is given by:

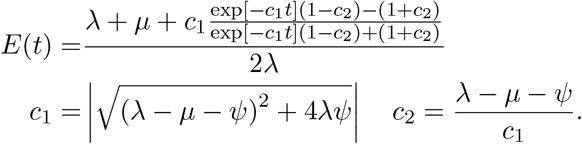
- Step 4: Derive *g*_*stem*_(*T*).

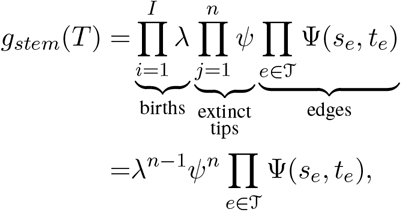

where *I* = *n* − 1.
- Step 5: Critical time representation. Given the assumption of constant rates and no Poisson sampling the expression for Φ(*t*) simplifies to:

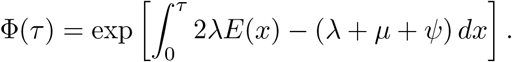 Hence we have:

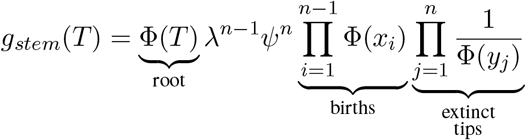
- Step 6: Impose conditioning 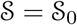.

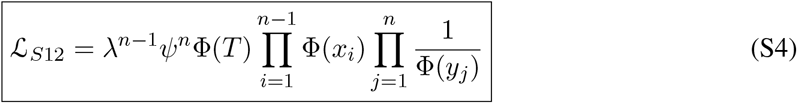

##### S1.6 Stadler *et al.* 2013 and Gavryushkina *et al.* 2014

Here we derive the tree likelihood given by **Theorem 1** in (Stadler and Bonhoeffer, 2013) and due to their shared the likelihood given by **Equation 4** in (Gavryushkina et al., 2014).

- Step 1: Specify the model.

– Piecewise constant Poissonian birth, death, and sampling rates.

* define transition times 0 = *t*_0_ *< t*_1_ < ... < *t*_*L*+1_ = *T*.
* *λ*(*τ*) = *λ*_*l*_*t*_*l*_ < *τ* ≤ *t*_*l*+1_.
* *μ*(*τ*) = *μ*_*l*_*t*_*l*_ < *τ* ≤ *t*_*l*+1_.
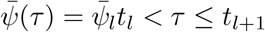.
– Fossils/Ancestral sampling:

* (Stadler and Bonhoeffer, 2013): No fossils *r* = 1.
* (Gavryushkina et al., 2014): piecewise-constant rate *r*(*τ*) = *r*_*l*_ *t*_*l*_ < *τ* ≤ *t*_*l*+1_.
– Concerted sampling attempts at each internal transition time *t*_*l*_ where *l* = 1, 2*...L*.

* The probability of a lineage being sampled during the CSA at time *t*_*l*_ is *ρ*_*l*_.
* The resulting total sampling rate is given by:

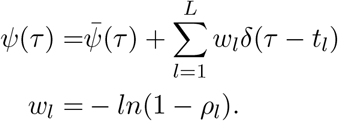
– Conditioning:

* Stadler et al. (Stadler and Bonhoeffer, 2013)—At least one sampled lineage, 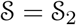.
* Gavryushkina et al. (Gavryushkina et al., 2014)— At least one sample 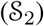 and a constant giving the number of un-oriented phylogenies 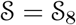.
- Step 2: Derive IVP for *g*_*e*_(*τ*). As in section 1 we have:

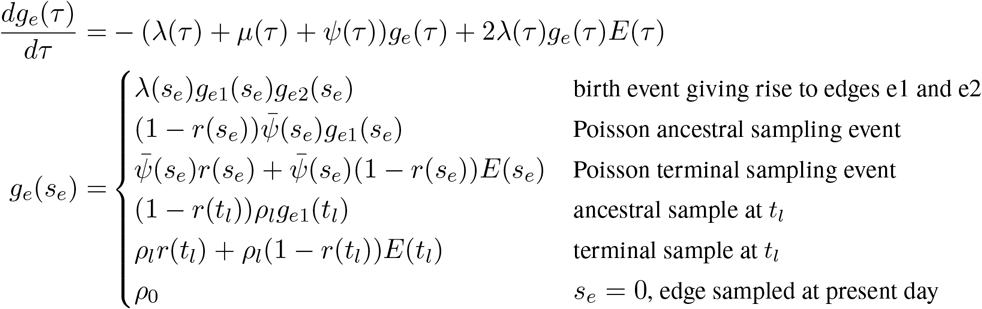
- Step 3: Derive IVP for *E*(*τ*). The IVP for *E*(*τ*) follows from Equation (8).

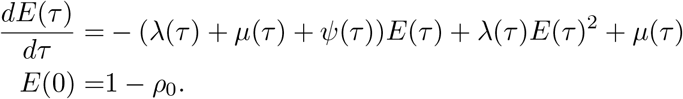 Given the piecewise constant nature there is a known general solution. Let *E*(*τ*) = *E*_*l*_(*τ*) where *t*_*l*_ < *τ* ≤ *t*_*l*+1_. Then define 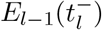 as the solution up to but not including the CSA at time *t*_*l*_ we have:

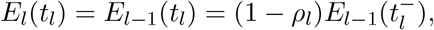

where *E*_*l*_(*τ*) is given by a solution similar to that in Equation (A1).

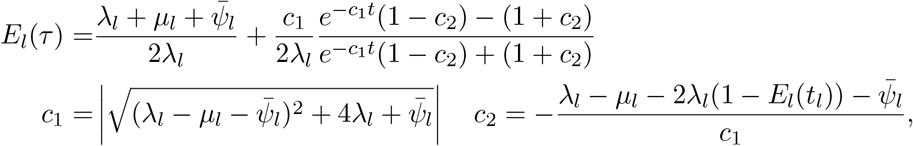

where *E*_0_(*t*_0_) = 1 − *ρ*_0_.
- Step 4: Derive *g*_*stem*_(*T*). As in section 1 the edge representation of *g*_*stem*_ is given by:

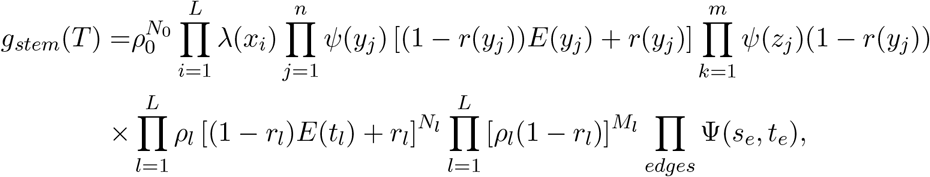

where Ψ(*s*_*e*_, *t*_*e*_) is given by Equation (6).
- Step 5: Critical time representation. We once again define the sub-flow Φ_*l*_(*τ*) where *t*_*l*_ < *τ* ≤ *t*_*l*+1_:

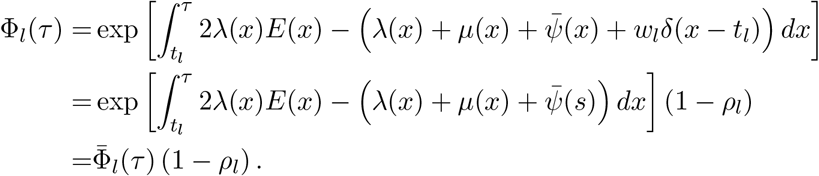 The complete flow is, as given by Equation (A3).

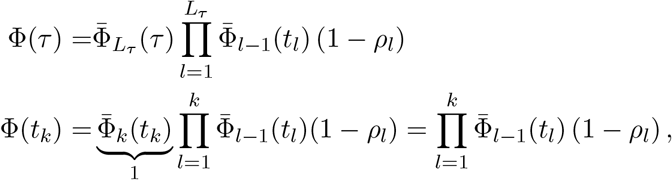

where *L*_*τ*_ is once again the largest index *l* such that *t*_*l*_ < *τ*. The critical time representation of *g*_*stem*_ from the previous stem gives the following.

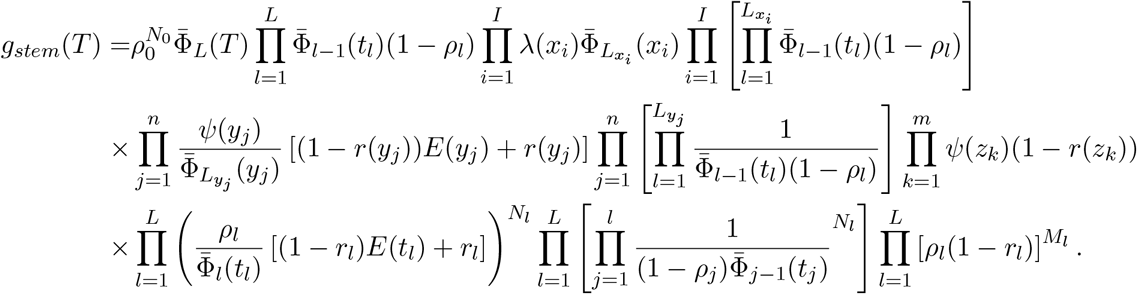 We can then simplify the likelihood using the relationships similar to those given by Equation (A4).

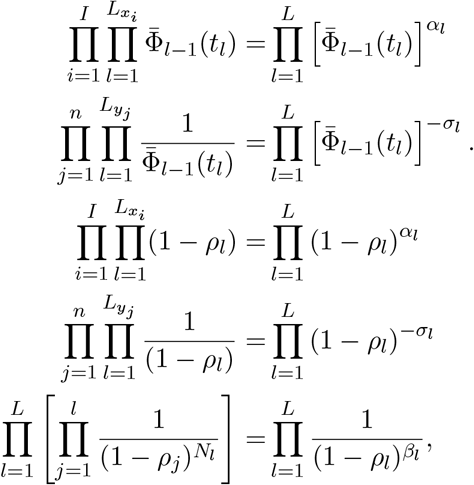

where *α*_*l*_ is the number of birth events before time *t*_*l*_ and *σ*_*l*_ is the number of Poisson sampling events before time *t*_*l*_ and *β*_*l*_ is the number of lineages sampled during the CSAs up to and including at time *t*_*l*_. The resulting simplified likelihood is given by:

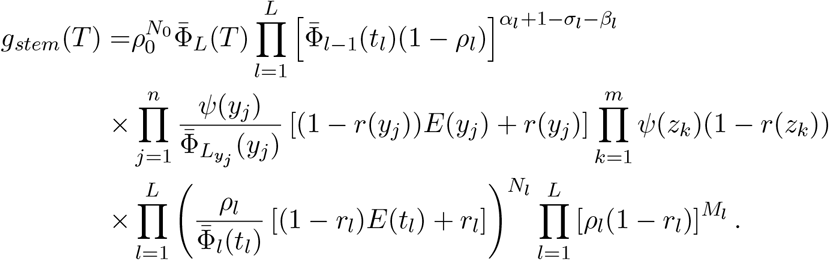
- Step 6: Conditioned likelihood.

– Stadler et al. (Stadler and Bonhoeffer, 2013) conditions on observing at least one sample. In addition the likelihood given by Stadler et al. assumes that all lineages are removed upon sampling *r*_*l*_ = 1 an assumption we will apply now. The likelihoods can be simplified by letting *n*_*l*_ = *α*_*l*_ + 1 − *σ*_*l*_ − *β*_*l*_ be the number of lineages that are alive immediately following the concerted sampling attempt at time *t*_*l*_

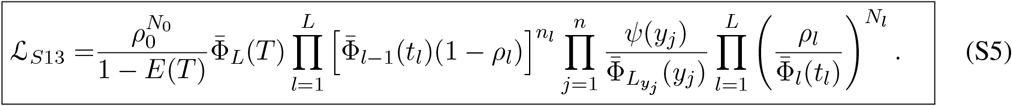
– Gavryushkina et al. (Gavryushkina et al., 2014) also condition on observing at least one lineage as well as multiply by a constant giving the number of un-oriented trees.

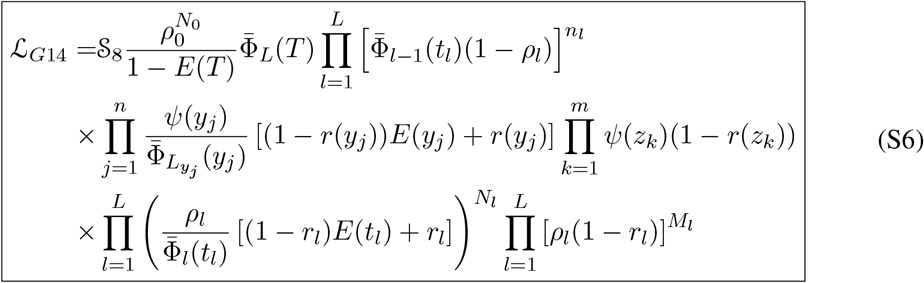

##### S1.7 Heath *et al.* 2014

- Step 1: Specify the model.

– Constant rates: *λ*(*τ*) = *λ*, *μ*(*τ*) = *μ*, *Ψ*(*τ*) = *Ψ*.

* Here *Ψ* denotes the sampling rate of fossils.
– Birth-death model with fossils, *r* = 0.
– Conditioned on observing ≥ 1 sample given *t*_*MRCA*_ 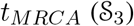 and enumerated over all possible attachments of fossils (terminal sampling events before the present day 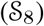).
- Step 2: Derive IVP for *g*_*e*_(*τ*.

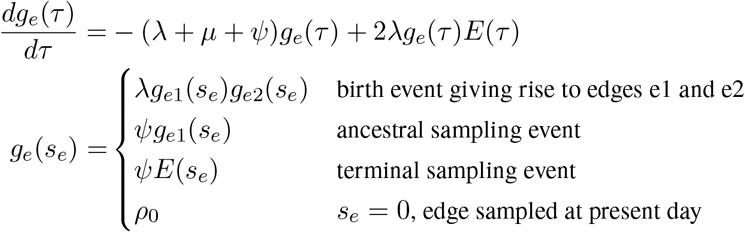
- Step 3: Derive IVP for *E*(*τ*.

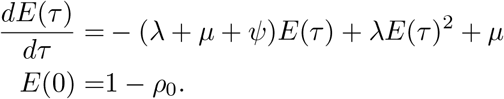 This has the same general solution as given above in section S1.2.
- Step 4: Derive *g*_*stem*_(*T*).

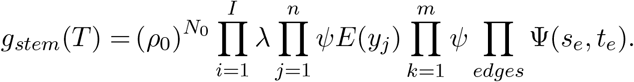
- Step 5: Critical time representation.

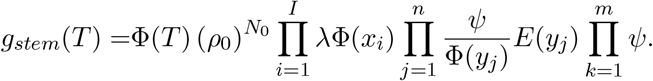
- Step 6: Conditioned likelihood. Conditioning on the probability that both daughter clades of the MRCA have at least one sampled extant descendant. 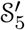. The general solution to *Ê*(*τ*) is known as given by the solution to *E*(*τ*) in section S1.1. Conditioning on all the possible attachments of the fossils (terminal samples before the present day). This requires multiplication by a constant 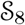. Let *γ*(*τ*) be the number of lineages alive at time *τ*. The the resulting number of fossil trees is.

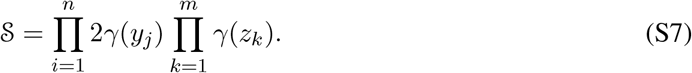

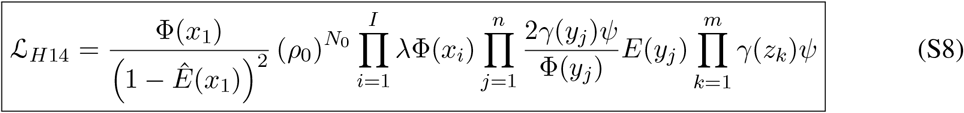

#### S2 Tables

##### General Birth-Death-Sampling Model Notation

**Table S1:**
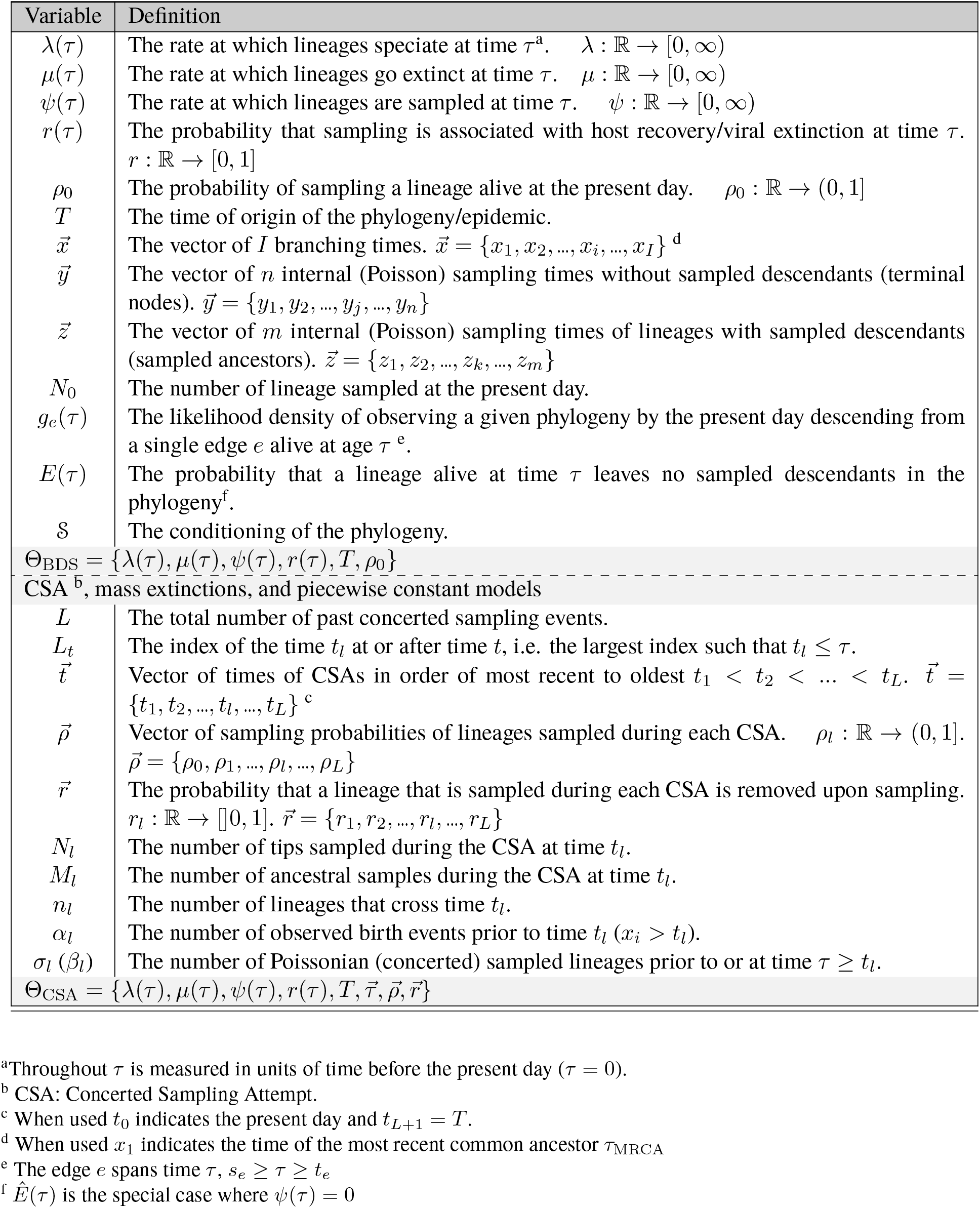
model notation.

**Table S2:** Relationship of single-type sub-models to the general BDS model.

| Model | $\lambda(\tau), \mu(\tau), \psi(\tau)$ | $r(\tau)$ | $\rho_0$ | $\rho_l$ | $r_l$ | $\nu_l$ | $\mathcal{S}$ | Section |
| --- | --- | --- | --- | --- | --- | --- | --- | --- |
| Stadler 2009 | constant, $\psi = 0$ | NA | $> 0$ | 0 | NA | 0 | $\mathcal{S}_6 \mathcal{S}_8$ | S1.1 |
| Stadler 2010 | constant | 0 | $> 0$ | 0 | NA | 0 | multiple | S1.2 |
| Morlon et al. 2011 | $\psi = 0$ | NA | $> 0$ | 0 | NA | 0 | $\mathcal{S}_1 = \mathcal{S}_2$ | S1.3 |
| Stadler 2011 | piecewise, $\psi = 0$ | NA | $> 0$ | 0 | NA | $\geq 0$ | $\mathcal{S}_3 = \mathcal{S}_5$ | S1.4 |
| Stadler et al. 2012 | constant | 1 | 0 | 0 | NA | 0 | $\mathcal{S}_1$ | S1.5 |
| Stadler and Bonhoeffer 2013 | piecewise | 1 | $> 0$ | $\geq 0$ | 1 | 0 | $\mathcal{S}_1$ | S1.6 |
| Gavryushkina et al. 2014 | piecewise | piecewise | $> 0$ | $\geq 0$ | $\geq 0$ | 0 | $\mathcal{S}_1 \times \mathcal{S}_7$ | S1.6 |
| Kühnert et al. 2014 | stochastic SIR | constant | 0 | 0 | NA | 0 | ?<sup>a</sup> |  |
| Heath et al. 2014 | constant | 0 | $> 0$ | 0 | NA | 0 | $\mathcal{S}_6$ | S1.7 |

<sup>a</sup> No specific conditioning mentioned.

**Table S3:**
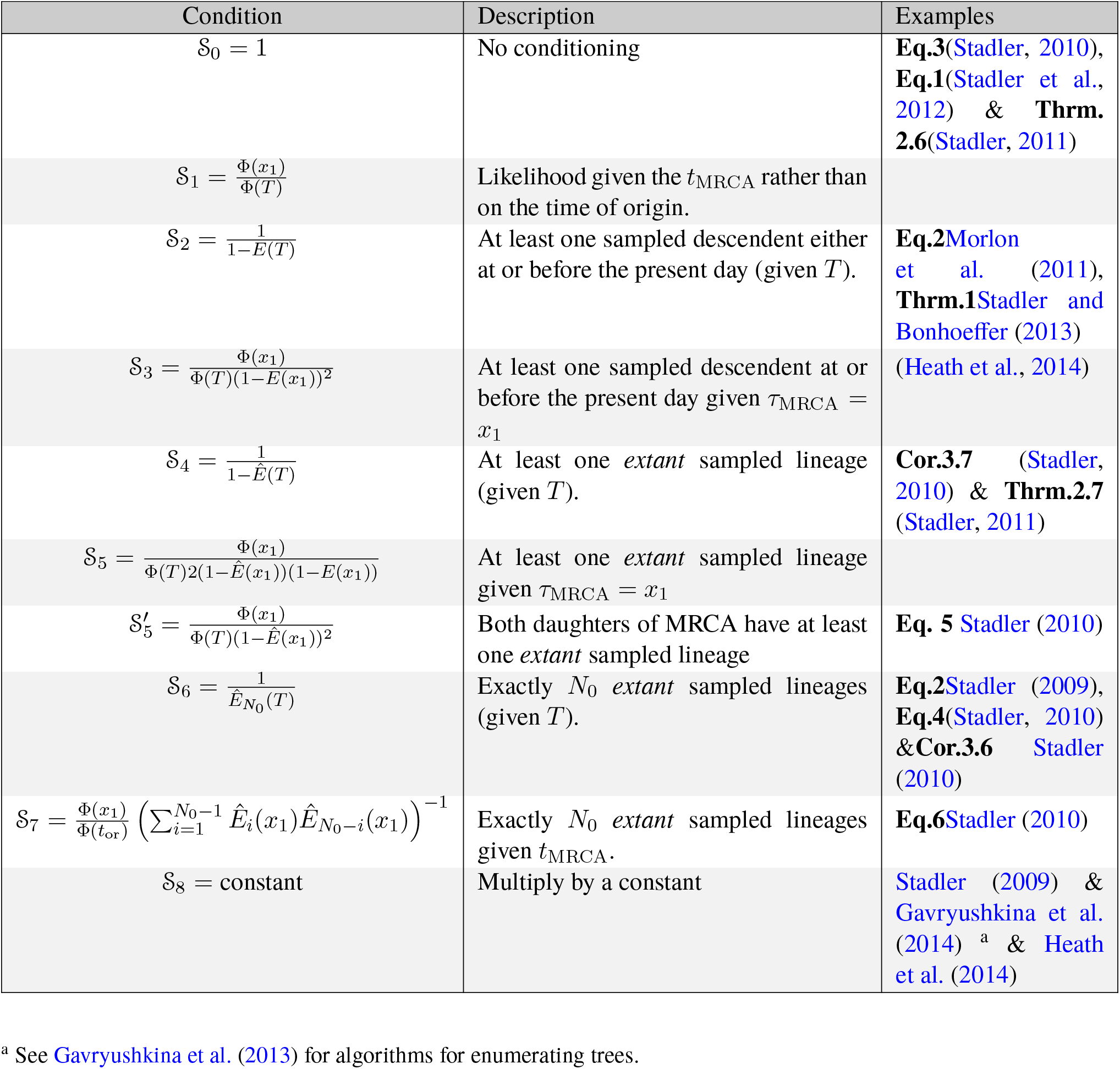
Alternative conditioning of tree likelihood.

## Part II

### The multi-type model

#### S3 The multi-type diversification model

Here we consider the diversification of lineages of *A* discrete types. Lineages of type *a* ∈ {1, 2*, ...A*} speciate/give birth to lineages of type *b ∈* {1, 2*, ...A*} at the time-variable rate *λ*_*a,b*_(*τ*) at time *τ* before the present day. As above we will use *τ* to denote time moving backward from the present day (*τ* = 0) to the origin of the phylogeny *τ* = *T*. When *a* = *b* speciation occurs without cladogenetic change, when *a* ≠ *b* specition is coincident with state change. In addition to the lineages changing state at birth events, lineages can mutate anagenetically from state *a* to state *b* at rate *γ*_*a,b*_(*τ*). Lineages of type *a* alive at time *τ*, go extinct/die at rate *μ*_*a*_(*τ*). For *τ >* 0 lineages are sampled at rate *Ψ*_*a*_(*τ*). Upon sampling a lineage may be removed from the population (e.g., sampling is coincident with treatment) with the state-dependent probability *r*_*a*_(*τ*). Finally, all lineages alive at the present day are sampled with a state-dependent probability *ρ*_*a*_. Model notation is summarized in Table S4.

As with the single-type model, the result of the mutli-type diversification process is a *full* and a *sampled* tree. Now however the tree is characterized by its topology 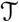 and the *colouring* of the tree 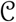 denoting the states of each lineage through time. Below we first derive an expression for the likelihood of a given coloured tree, 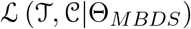. However, as plausibly attainable data consists of knowledge at only some or all the sampled ancestral nodes and/or tips, we must then integrate the likelihood 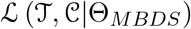 over all possible tree colourings consistent with the observed data.

##### S3.1 Derivation of 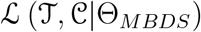

We derive the likelihood using the steps used above for the single-type birth death sampling model. As with the single-type model for the likelihood calculation be begin by representing the phylogeny as a series of edges. However, as we are now referring to the coloured phylogeny we consider the set of all coloured edges, with each edge being a a segment of the phylogeny that is all of one colour beginning and ending at birth, sampling, or mutation events. Specifically, moving backward in time towards the tree origin let edge *e* start at time *s_e_* before the present at a birth event, sampling (ancestral or tip), or mutation event, and continue toward the tree origin until time *τ*_*e*_ ending at either a birth event, ancestral sampling event, mutation event or at the tree origin.

As with the single-type model the tree likelihood depends on two different functions. First, *g*_*e,a*_(*τ*) is the probability that an edge *e* with state *a* alive at time *τ* (hence *t*_*e*_ > *τ* > *s*_*e*_) gives rise to the subsequently observed phylogeny between *τ* and the present day. Second, *E*_*a*_(*τ*) is the probability a lineage of type *a* alive at time *τ* has no sampled descendants between *τ* and the present day. We begin below by first deriving the initial value problems for these two functions.

##### Step 1: Derive the Initial Value Problem for *g*_*e,a*_(*τ*)

To simplify notation we define the total birth and mutation rates of a given type Λ_*a*_ = ∑_*b*_λ_*a,b*_ and Γ_*a*_ = ∑_*b*_λ_*a,b*_ and the relative probability a birth or mutation event was of a given type, 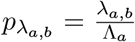 and 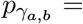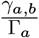. For some small amount of time Δ*τ* the recursion equation for *g*_*e,a*_(*τ*) is given by:

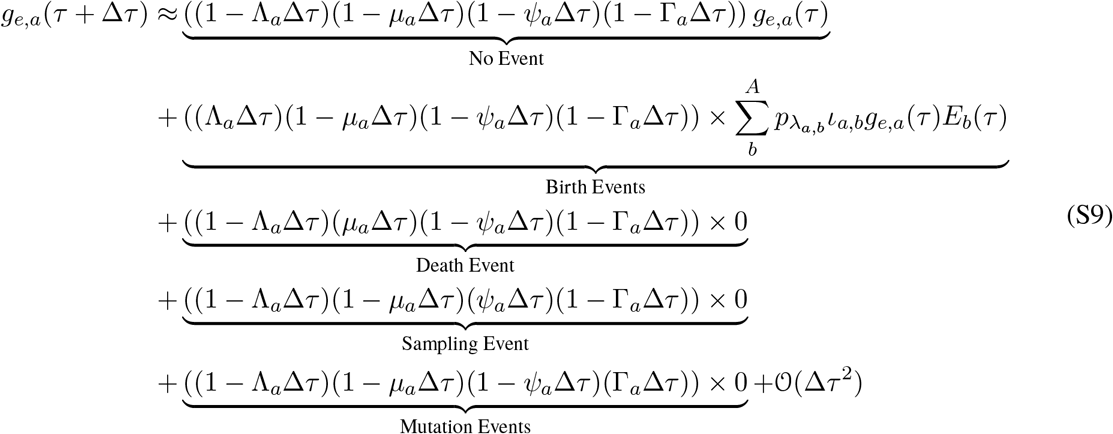

where *ι*_*a,b*_ is an indicator variable that has value 2 when *a* = *b* and 1 otherwise. This recursion equation uses the fact that edges are all in state with mutation events resulting in the end of an edge and the origin of a daughter edge with a different state. This is reflected by the initial conditions of *g*_*e,a*_(*τ*) as given below in equation (S11).

As above we can then use the definition of a derivative to obtain the corresponding differential equation

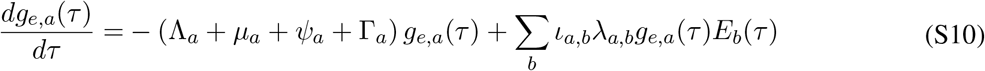

The initial conditions for *g*_*e,a*_ at the start of the edge, time *s*_*e*_, depend on the event that occurs in the sampled tree at that time. Specifically there are five types of events in the observed tree. 1) *Observed birth events* where a lineage of type *a* speciates producing a new lineage of type *b*. 2) *Ancestral sampling events* where a lineage of type *a* is sampled, is not removed from the population, and then has subsequently observed descendants. 3) *Terminal sampling events* where a lineage of type *a* is sampled and then either is immediately removed from the population or remains in the population but has no sampled descendants. 4) *Transition events* where a lineage of type *a* transitions to a lineage of type *b*. There are two possible ways a transition event can occur. First, a lineage can switch states due to a anagenetic *mutation event* or second an apparent transition can arise due to a birth event with cladogenetic state change followed by subsequent extinction of the parental lineage. We refer to this second form of transition event as a *hidden birth event*.

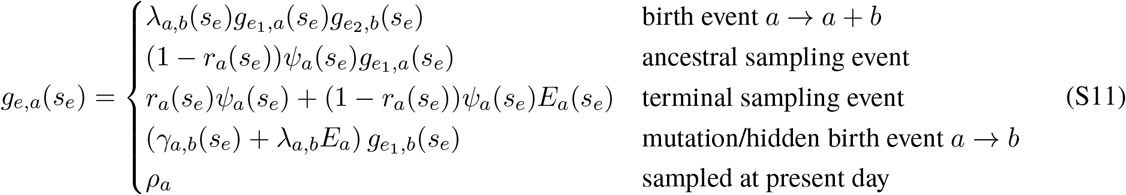

Importantly, as the ODES in equation (S10) are linear the corresponding IVP has the following general solution given the initial conditions above.

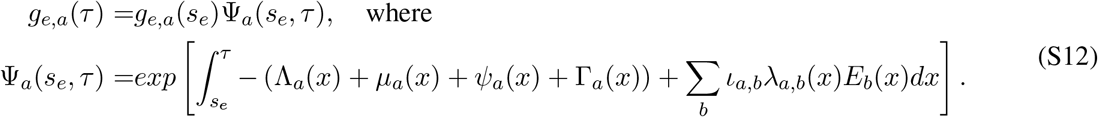

The IVP for the function *E*_*a*_(*τ*) can be derived in an analogous manner.

##### Step 2: Derive the Initial Value Problem for *E*_*a*_(*τ*)

Once again we begin by deriving a recursion equation for the change in *E*_*a*_(*τ*) over from time *τ* to *τ* + Δ*τ*.

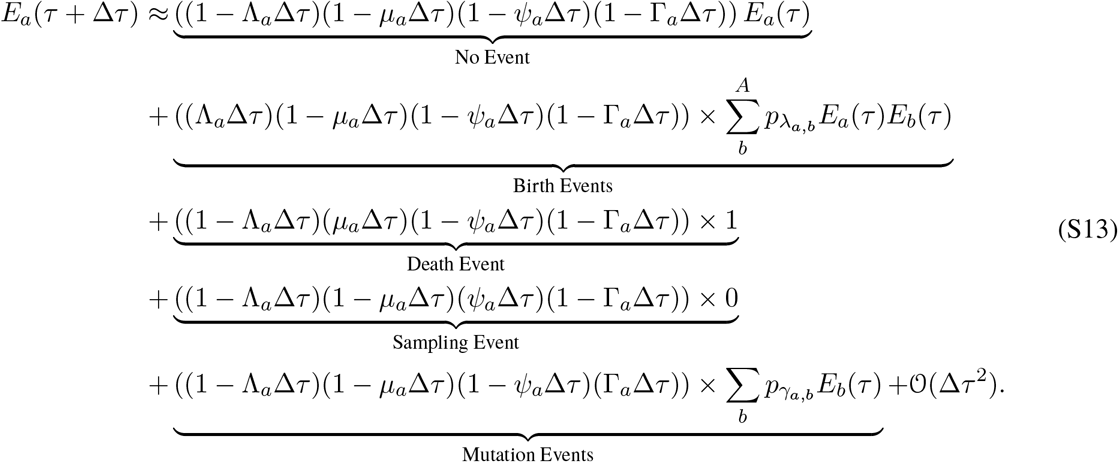

Using the definition of a derivative we have:

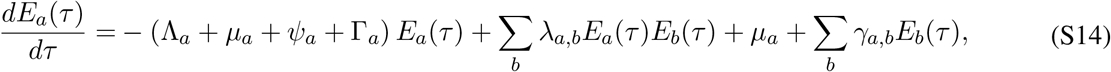

with initial conditions given at the present day by:

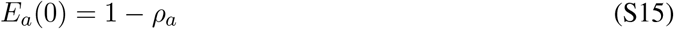

##### Step 3: Derive Expression for *g*_*stem*_(*T*)

As in equation (9) for the single-type model, the tree likelihood is given by the value of the function *g* of the stem edge evaluated at the time of the origin of the phylogeny, *T*. From the solution to the initial value problem given by equation (S12) and the initial conditions (S11) we can write *g*_*stem*_(*T*) as a product over the events in the tree and all the probability flow Ψ of the edges in-between. To do so let 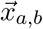 be a vector of length *I*_*a,b*_, giving the time before the present day of all the observed birth events where lineages of type *a* give rise to lineages of type *b*. Let 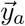 be a vector of length *J*_*a*_ giving the sampling time of tips of type *a* and 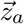 a vector of length *K*_*a*_ the sampling type of sampled ancestors of type *a*. Finally let 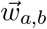 be the time of observed transition events where a edge of type *a* transitions becomes an edge of type *b*. Once again, transition events can arise due to both direct mutation and hidden birth events. The resulting expression for the likelihood *g*_*stem*_(*T*) is given by:

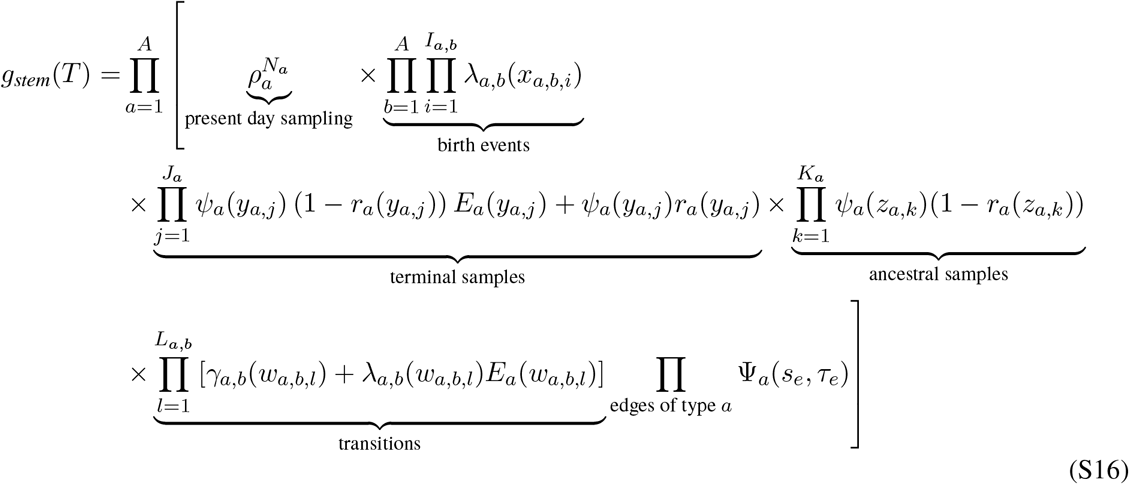

##### Step 4: Rewrite *g*_*stem*_(*T*) in Terms of Critical Times

Rather than enumerate Ψ over the edges of the phylogeny we can rewrite equation (S16) in terms of only the critical times 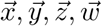 Written in this form the likelihood also depends on *c*\* the colour of the phylogeny at the origin. To do so we define Φ_*a*_(*τ*) = Ψ_*a*_(0*, τ*) and rewrite the probability flow Ψ_*a*_(*s*_*e*_, *τ*) as a ratio:

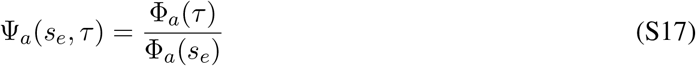

Substitution into equation (S16).

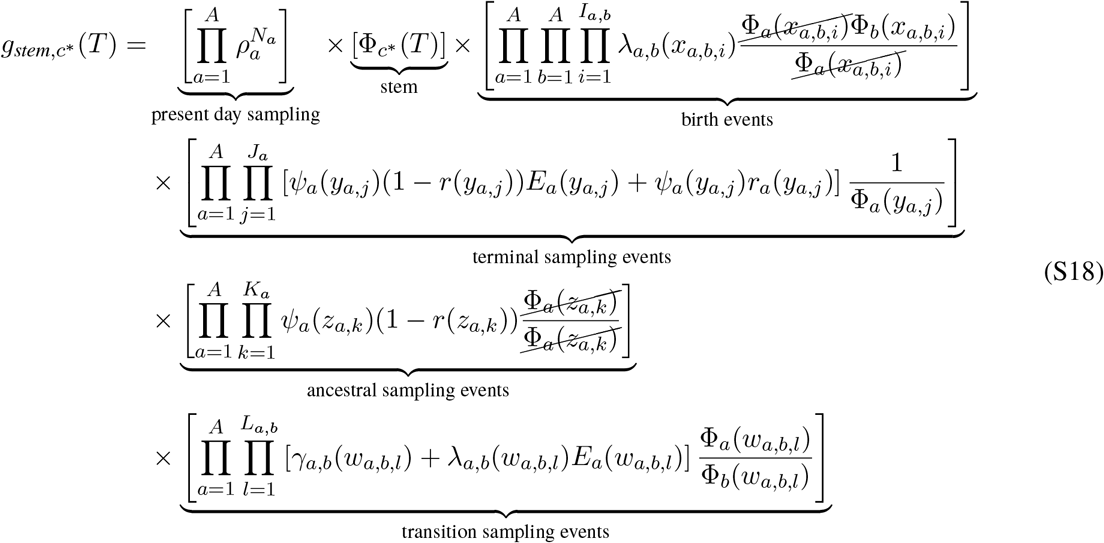

##### Step 5: Condition the Likelihood

We conclude, as in the single type model by multiplying the likelihood by including a general form of conditioning 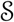.

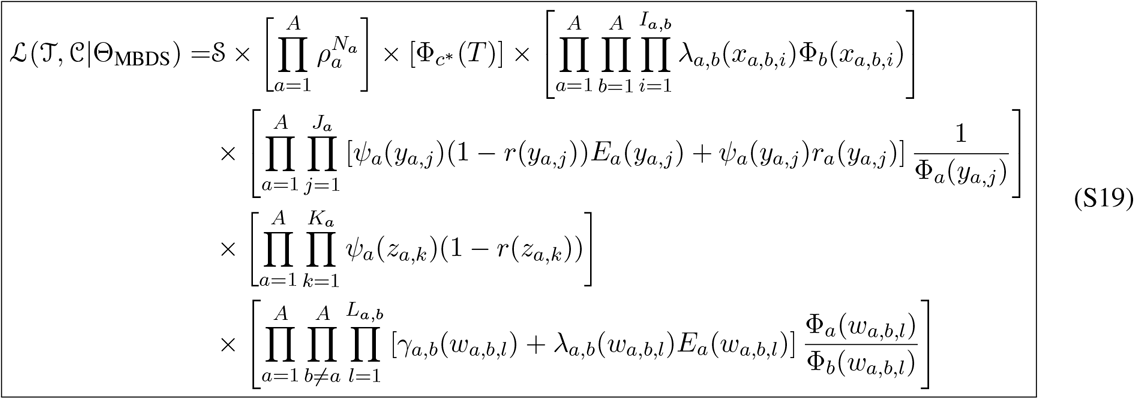

Due to the combination of cladogenetic change and different possibility of hidden birth events due to extinction and sampling, there are six different types of birth events included in the likelihood of the sampled tree. The following diagram summarizes these different birth events and how each is included in the equation (S19).

**Figure S1:**
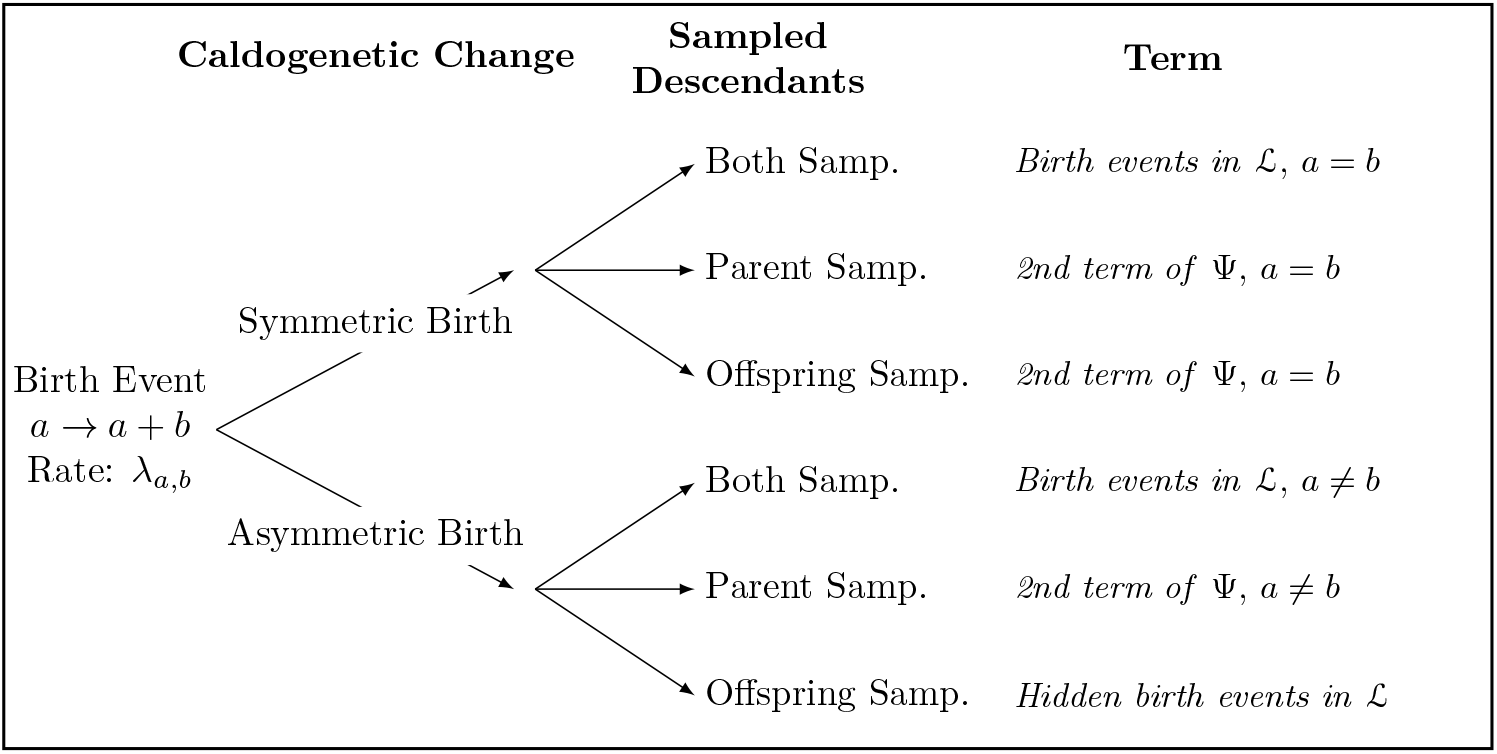
The six possible birth events and how each in included in the likelihood given by equation (S19).

##### S3.2 Integrating over tree colourings

The likelihood expression given by equation (S19) requires knowledge of when and how (e.g, through clado-genetic or anagenetic change) lineages have changed state, information which it is not plausible to obtain. Instead the available data is likely to include either only knowledge of the sampled tip states, which we denote as 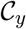 for the colouring at the points in 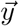, and potentially the known states of sampled ancestors, 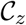. Note that the 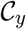 and 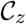 need not include information for all sampled tips or sampled ancestors respectively. Calculating the corresponding likelihood of of the tree topology and the observed data can be calculating by computing the following integral:

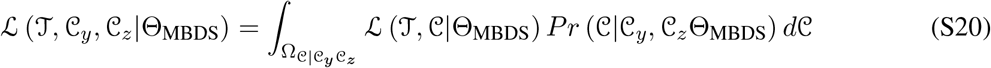

where 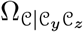 is the space of all possible tree colourings that are consistent with the observed states at the sampled tips and sampled ancestors. Here 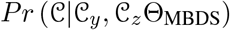 is the probability of observing a given tree colouring given the observed states. Numerical calculation of this integral can be obtained via Monte Carlo through the use of a stochastic mapping algorithms (Nielsen, 2002) as exemplified below.

#### The BiSSE-ness model with time-variable rates

Here we examine one particular sub-model of the more general multi-type model which exemplifies many of the complexities of diversification with multiple states. Specifically we extend the BiSSEness/ClaSSE model (Magnuson-Ford and Otto, 2012; Goldberg and Igić, 2012) to allow for time-variable model rates. See *Mathematica* notebook for reference

##### The diversification model

The multi-type model above allows the birth, death, and sampling rates to vary arbitrarily through time. In order to specify realistic time-variable rates we begin by specifying a mechanistic model for the diversification process. Specifically we model the diversification dynamics with density-dependence where birth and death rates vary respect to a model of deterministic logistic growth with mutation. The deterministic number of lineages of type *a* through time, *n*_*a*_(*t*), is given by the solution to the following initial value problem:

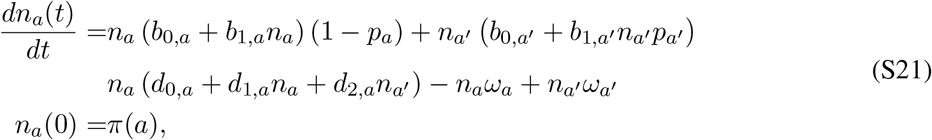

where *p*_*a*_ is the probability that a birth event with a type *a* parental lineage undergoes cladogenetic change and *π*_*a*_ is the prior probability the tree is initially in state *a*. The mutation/transition rate from state *a* to *a′* is constant, occurring at a per-capita rate *ω*_*a*_. Under this mechanistic model the diversification model rates are given by:

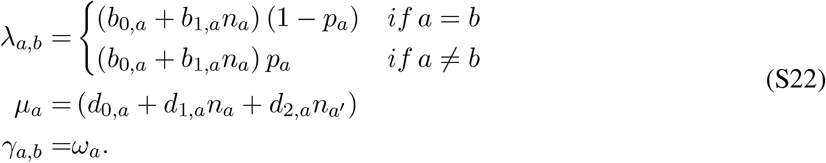

Lineages of type *a* are sampled at the present day with probability *ρ*_*a*_.

##### The BiSSEness likelihood with time-variable rates

The likelihood for the tree given the observed tip states can be derived in a similar manner as in (Magnuson-Ford and Otto, 2012), allowing the birth and death rates to vary through time.

First we derive the initial value problem for *D*_*e,a*_(*τ*), the probability an edge *e* in the tree topology 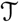 in state *a* at time *τ* gives rise to the subsequently observed phylogeny. Note that in contrast to the function *g*_*e,a*_(*τ*), the edge *e* here refers to an edge in the tree topology not in a coloured tree and hence can change state. The edge here begins at either the origin of the phylogeny or a birth event and ends at a birth event or at the tip in the phylogeny.

We begin by deriving the recursion equation for *D* between time *τ* and *τ* + Δ*τ*.

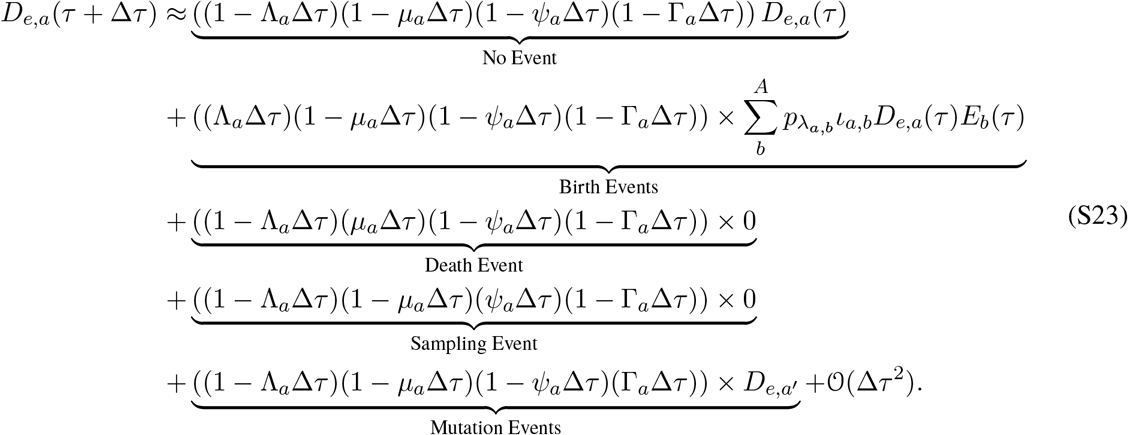

This equation is similar to that for *g*_*e,a*_ except for the mutation term which is no-longer multiplied by 0, but rather by *D*_*e,a′*_. As a result the corresponding IVP is a system of coupled differential equations and no-longer has a known general solution. Similarly, as the type of birth event occurring at each internal node is now unknown, we must sum over the probability of all possible events at these nodes.

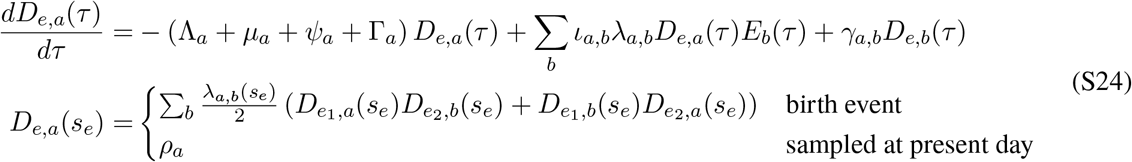

The initial value problem for *E*_*a*_(*τ*) is identical to that given in equation (S14) and (S15). The likelihood of the tree is given by the sum of the probability *D*_*e,a*_ across states *a* weighted by the prior probility of those states.

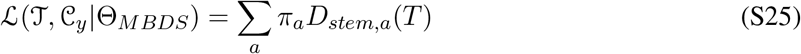

##### Stochastic mapping and integration over colourings

The likelihood of a given coloured tree is given by equation (S19). To calculate the likelihood of the tree where only the tip states are known as in the BiSSEness likelihood above, we must integrate over all possible colourings of the tree as specified in equation (S20). Such an integral can be performed by Monte Carlo integration, first simulating tree colorings consistent with the observed tip states from the distribution 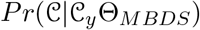. By then taking the mean of the likelihoods of each of these coloured trees we can approximate the tree likleihood with known tip states 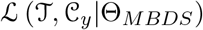. To sample such colourings we using a stochastic extend the stochastic mapping method first developed by (Nielsen, 2002) allowing for time-varying transition rates and cladogenic state change.

##### Preliminaries

- Let *Q*(*t*) be the transition rate matrix at time *t* such that *Q*_*j,i*_(*t*) is the rate at which lineages in state *i* (column) transitions to state *j* (row). For the multi-type diversification process this transition rate is given by:

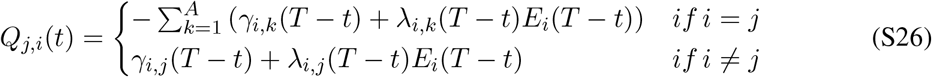 Note that the matrix *Q* is defined forward in time from the origin to the present whereas the diversification rates are defined backward in time. To allow for the computation of the transition probability matrix *P* defined below, we approximate the time-variable transition rates with a piecewise constant function such that for the interval (*l* − 1) * Δ*t < t < l*Δ*t* the transition rate matrix is assumed to be constant as given by:

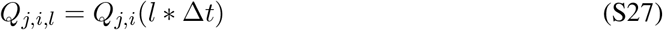

where 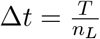 is a small uniform time and *l* ∈ {1, 2*, ...n*_*L*_}.
- The transition probability *P*_*j,i,l*_ of transitioning from state *i* to state *j* between (*l* − 1) * Δ and Δ*t* can be approximated as:

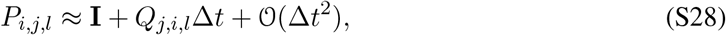

where **I** is the identity matrix.
- Let *P*_*j,i*_(*t*_0_*, t*_*F*_) be the probability of transitioning from state i at time *t*_0_ to state j at time *t*_*F*_. Define *L*(*t*) as the integer index such that (*L*(*t*) − 1)Δ*t < t < L*(*t*)Δ*t*.

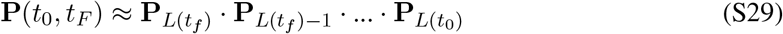 Assuming Δ*t* is small this approximation assumes that no events occur between times *t*_0_ and *L*(*t*_0_)Δ*t* and between *t*_*F*_ and *L*(*t*_*F*_)Δ*t*.
- Let *D*_*K*_ be the tip states of the descendants of node *K*.
- Let *Y*_*K*_ bet the state of node *K*.
- With respect to stochastic mapping an edge refers to a *topological edge*, spanning between two nodes in the tree topology 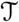. This is in contrast to an edge in the coloured tree, as used in equation (S16), which is a portion of topological edge that is all one colour/state.

**Figure S2:**
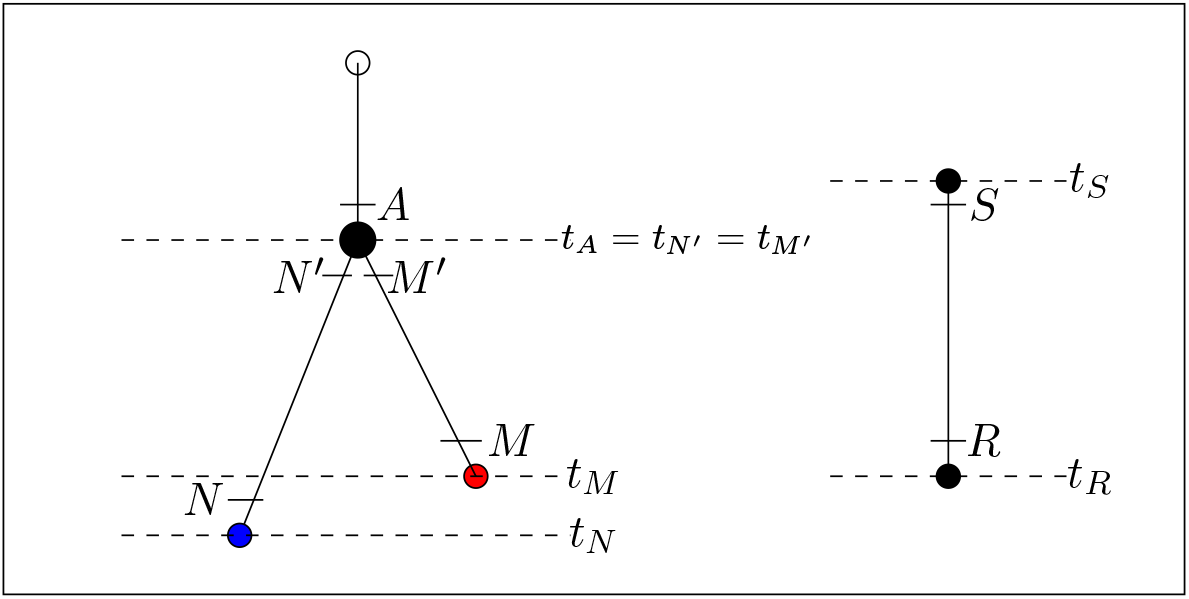
The two edge motifs in the tree with respect to stochastic mapping: A) a branching event where an edge *A* gives rise to a edge *N* and an edge *M* and B) the stem edge between the *stem node* at the tree origin *S* and the *root node* at the most recent common ancestor *R*.

##### Step 1: Calculate *f*_*K,i*_ = *Pr*(*D*_*K*_|*Y*_*K*_ = *i*)

Define *f*_*K,i*_ = *Pr*(*D*_*K*_|*Y*_*K*_ = *i*), the probability of observing the known descendants of node *K*, *D*_*K*_, given that node *K* is in state *i*.

**For each tip node** *T*:

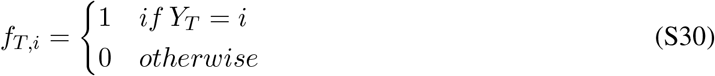

**For each “originating” node** (e.g, *N′* and *M′* in Figure S2):

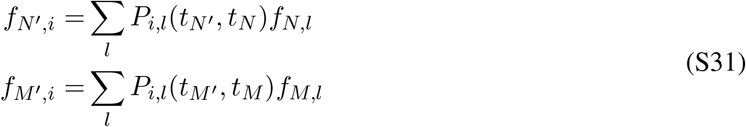

**For each “ancestral”/“ending” node** (e.g., *A* in Figure S2)

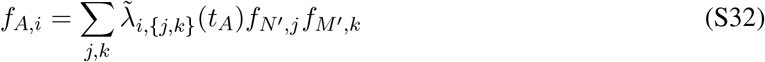

where 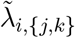 is the probability that a lineage of type *i* gives rise to a linage of type j and a linage of type k given that a birth event occurs.

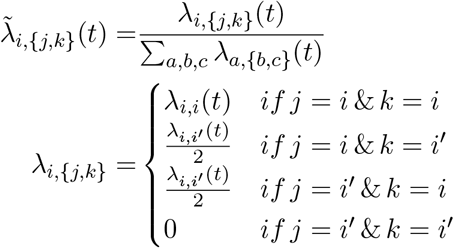

Using Felsenstein’s pruning algorithm, equations (S30), (S31), and (S32) can be used to calculate the probability *f*_*K,i*_ for each node *K* and each state *i* in the tree.

##### Step 2: Calculate *Pr*(*Y*_*K*_ = *j*|*D*_*K*_, *Y*_*K*+1_ = *i*)+ simulate nodes, births

Let node *K* + 1 be the direct ancestor of node *K*.

**For the “stem” node:**

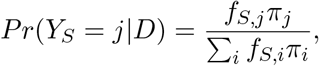

where *π*_*j*_ = *Pr*(*Y*_*S*_ = *j*) is the probability the stem node has state *j*, hence this result follows from Bayes theorem.

*Choose stem node state!.*

**For each “ending” node:**

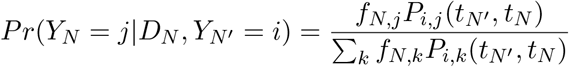

The *f*_*N,\**_ terms account for *Pr*(*Y*_*N*_ = *j*|*D*_*N*_) whereas the *P*_*i,\**_ terms give the *Pr(Y*_*N*_=*j*|*Y*_*N′*_=*i*).

*Choose ending node state!.*

**For birth event and ’originating’ node:** Calculate the probability of the different birth events

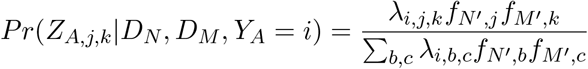

*Choose birth event!. Gives value of j and k.*

##### Step 3: Simulate the colourings of each edge

Given the originating state of each node we can then us a Gillespie algorithm with the transition matrix *P* to simulate the colouring along each branch. Edge colourings are simulated until one is generated that matches the simulated ending state chosen in step 2. From the simulated edges and nodes we can obtain the vectors 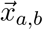 and 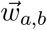 as well as the initial state of the tree *c*\* and the states of the tip nodes as summarized by *N*_*a*_.

**Figure S3:**
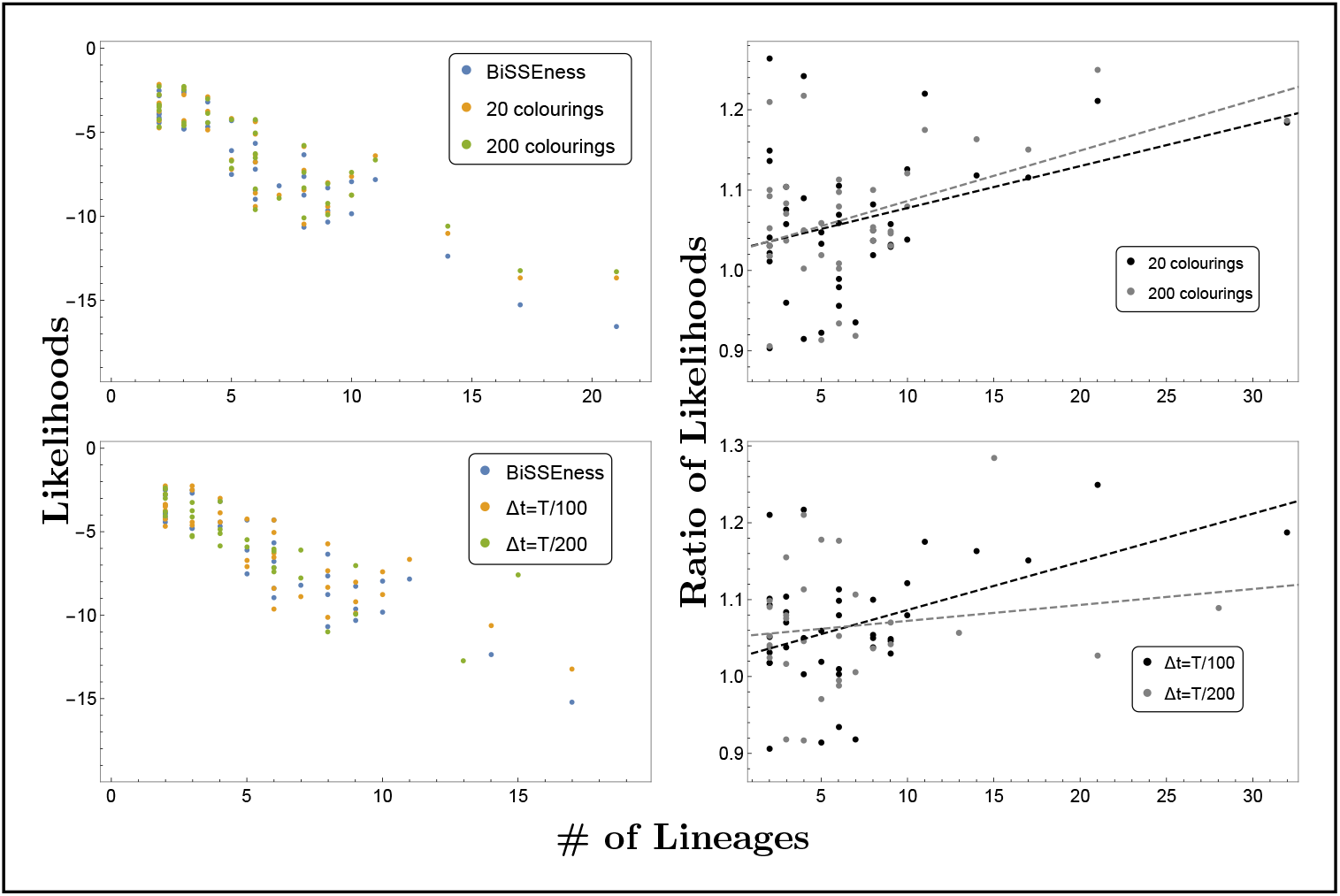
Comparing likelihoods. Comparing the tree likelihood calculated via Monte Carlo integration (see equation (S20)) (orange and green points) to likelihood calculated via direct numerical integration (equation (S25)). The error given by the ratio of the likelihoods as shown by the ratio of likelihoods in shown in the right hand plots. The accuracy of the likelihood is a foremost a function of tree size. The likelihood is reasonably accurate when relatively few terms are used in the Mote Carlo integration, e.g., averaging over only 20 colourings, and is improved by increasing the accuracy of the piecewise-constant approximation to the diversification rates (e.g., 200 vs. 100 piecewise-constant steps). Top row: Δ*t* = *T/*100, bottom row: 200 colourings.

#### S4 Tables

**Table S4:** Notation for the general multi-type model. Throughout *t* represents time moving forward from the origin (*t* = 0) of the tree to the tips (*t* = *T*) and *τ* represents time moving backward from the tips (*τ* = 0) to the origin (*τ* = *T*). The indices *a, b* denote lineage types and can take on integer values between 1 and *A*.

| Variable | Definition |
| --- | --- |
| $A$ | The number of discrete lineage types. |
| $\lambda_{a,\{b,c\}}(\tau)$ | The time-varying rate at which lineage of type $a$ gives rise to daughter lineages of type $b$ and $c$ at time $\tau$. |
| $\mu_a(\tau)$ | The time varying rate at which lineages of type $a$ go extinct at time $\tau$ |
| $\psi_a(\tau)$ | The time varying rate at which lineages of type $a$ are sampled. |
| $r_a(\tau)$ | The time varying probability that upon sampling lineages are removed. |
| $\gamma_{a,b}(\tau)$ | The time varying rate at which lineages of type $a$ transition to type $b$. |
| $\rho_a$ | The probability a lineage of type $a$ alive at the present day is sampled. |
| $N_a$ | The number of lineages of type $a$ sampled at the present day. |
| $\vec{x}_{a,b,c}$ | A vector of length $I_{a,b}$ giving the timing of speciation events where a lineage of type $a$ gives rise to a daughter of type $b$. |
| $\vec{y}_a$ | A vector of length $J_a$ giving the times at which lineages of type $a$ are sampled for which there are no sampled descendants (sampled tips). |
| $\vec{z}_a$ | A vector of length $K_a$ giving the times at which lineages of type $a$ are sampled for which there *are* sampled descendants (sampled ancestors). |
| $\vec{w}_{a,b}$ | A vector of length $L_{a,b}$ giving the times at which lineages of type $a$ mutate to lineages of type $b$. |
| $c$ | The state at the origin (stem node) of the phylogeny. |
| $\mathcal{T}$ | The tree topology. |
| $\mathcal{C}$ | The colouring of the tree. |
| $g_e(a, \tau)$ | The probability an edge $e$ of type $a$ alive at time $\tau$ gives rise to the subsequently observed phylogeny |
| $E_a(\tau)$ | The probability a lineage of type $a$ alive at time $\tau$ has no sampled descendants between time $\tau$ and the present day. |
| $\iota_{a,b}$ | An indicator variable with value 2 if $a = b$ and 1 otherwise. |
| $\mathcal{C}_y$ | The known tip states |
| $\mathcal{C}_z$ | The known states of sampled ancestors |
| $\Theta_{\text{MBDS}}$ | The parameters of the multi-type BDS model |
| SSE Algorithm |  |
| $\pi_c$ | The probability the stem node has state $c$ |
| $D_{N,a}(\tau)$ | The probability a “topological” edge $N$ of type $a$ at time $\tau$ gives rise to the subsequently observed phylogeny. |

## Notes

### Competing Interest Statement

The authors have declared no competing interest.

### Summary of Updates

Updated Manuscript with Multi-Type Model.

